# The unique three-dimensional arrangement of Macrophage Galactose Lectin enables *E. coli* LipoPolySaccharides recognition through two distinct interfaces

**DOI:** 10.1101/2023.03.02.530591

**Authors:** Massilia Abbas, Meriem Maalej, Ferran Nieto-Fabregat, Michel Thépaut, Jean-Philippe Kleman, Isabel Ayala, Antonio Molinaro, Jean-Pierre Simorre, Roberta Marchetti, Franck Fieschi, Cedric Laguri

## Abstract

LipoPolySaccharides are a hallmark of Gram-negative bacteria and their presence at the cell surface is key for bacterial integrity. As surface exposed components, they are recognized by immunity C-type lectin receptors present on Antigen Presenting Cells. Human Macrophage Galactose Lectin binds *E. coli* surface that presents a specific glycan motif. Nevertheless, this high affinity interaction occurs regardless of the integrity of its canonical calcium-dependent glycan binding site. Nuclear Magnetic Resonance of MGL carbohydrate recognition domain and complete extracellular domain revealed a new glycan binding site opposite to the canonical site. A model of trimeric Macrophage Galactose Lectin was determined based on a combination of Small Angle X-ray scattering and Alphafold. A disulphide bond positions the Carbohydrate Recognition Domain perpendicular to the coiled-coil domain. This unique configuration for a C-type lectin orients the six glycan sites of MGL in an ideal position to bind LipoPolySaccharides at the bacterial surface with high avidity.

## INTRODUCTION

The outer membrane of Gram-negative bacteria is compositionally asymmetric with LipoPolySaccharides (LPS) covering most of its surface (Fig 1A), while phospholipids compose the inner leaflet. LPSs form a highly impermeable barrier and are critical in bacterial virulence (di Lorenzo et al., 2022); their structural variability and tight assembly protect bacteria against uptake of antimicrobials and enable evasion from host defenses. Constant transport and maintenance of LPS in the outer membrane is critical in the survival of bacteria. LPSs are composed of three moieties, the lipid A formed by N- and O-acylated di-glucosamine, the core OligoSaccharide (core OS) and O-antigen polysaccharide repeat (Fig 1B). These complex glycolipids are detected by the immune system through the lipid A *via* the well described LBP-MD2-TLR4 cascade (Ryu et al., 2017) and by the caspase system in the cytoplasm (Yi, 2017). Antibodies directed against the glycan moieties, core OS (Reinhardt et al., 2015) and O-antigen polysaccharides are also produced by the immune system to modulate bacterial infections (Rollenske et al., 2018). Another protein family present on Antigen Presenting Cells, C-type lectin receptors (CLRs), have been shown to bind sugars from the core OS of LPS (Geissner et al., 2019; Hanske et al., 2017; Jégouzo et al., 2020). CLRs are key immunity receptors which recognize a plethora of pathogen glycans (Mayer et al., 2017) and the interaction of these CLRs with their ligands, discriminating non-self from self-molecular motifs, allows dendritic cells to modulate the immune response towards either activation or tolerance (Mnich et al., 2020). Macrophage Galactose-type Lectin (MGL) is a trimeric type II CLR expressed on the cell surface of macrophages and dendritic cells (Fig 1D). It mediates interactions between endothelial and cancer cells (Bulteau et al., 2022) but also recognizes microbial glycans. Its main role appears to be an immunomodulatory activity, reducing excessive inflammatory responses. So far MGL has been described to recognize *S. aureus, C. jejuni, K. pneumonia, N. gonorrhea, B. pertussis and M. tuberculosis* (Mnich et al., 2019; Naqvi et al., 2021; van Sorge et al., 2009; Vliet et al., 2009).

**Figure 1:**
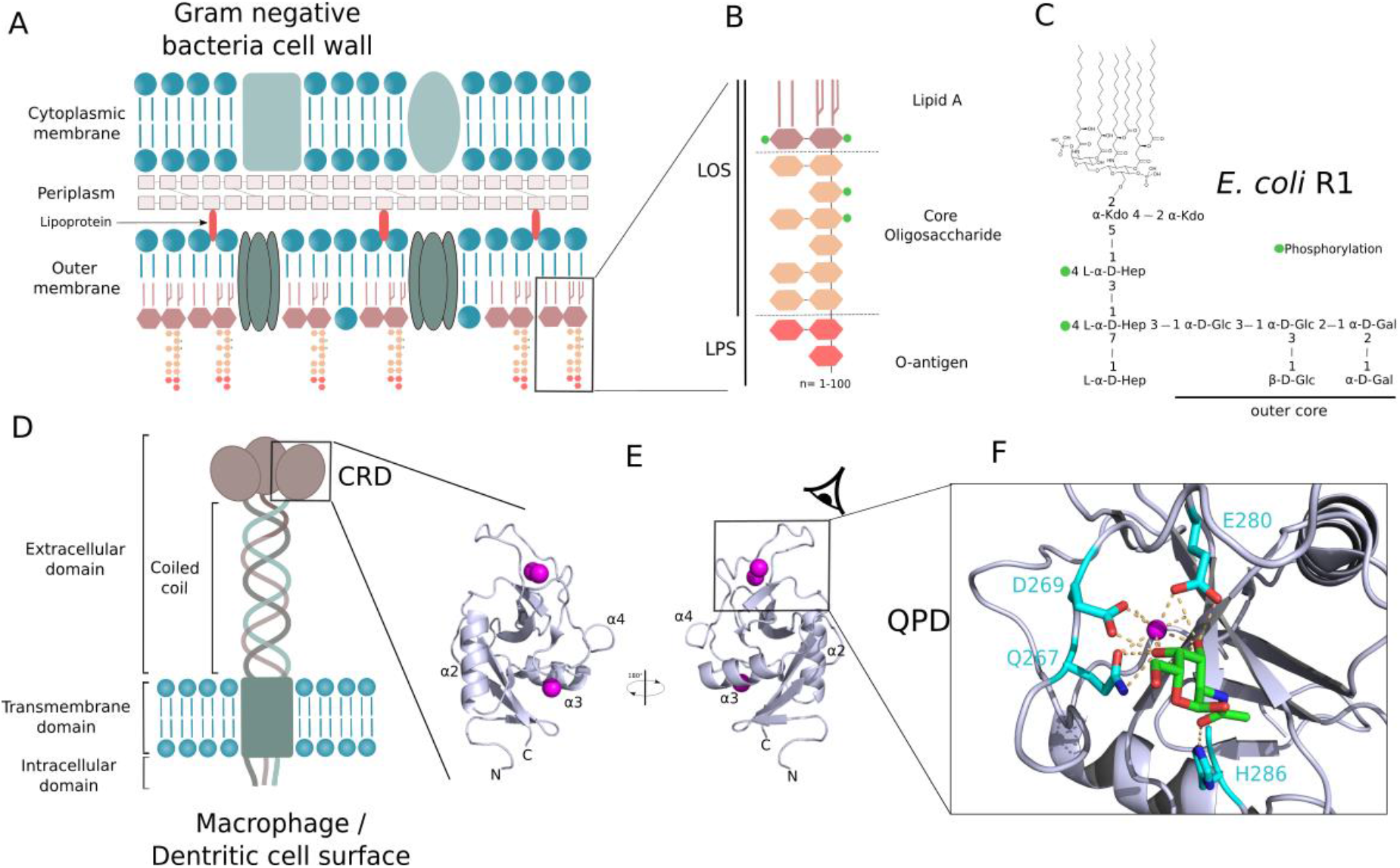
Organization of Gram-negative bacteria cell wall and of MGL. A) General structure of Gram-negative bacteria cell wall. B) LPS composing the outer leaflet of the outer membrane. C) Structure of *E. coli* R1 LipoOligoSaccharide (LOS) mainly used in this study. D) Domain organisation of MGL at the surface of Antigen Presenting Cells. E) Structure of CRD domain of MGL. F) Close up view on GalNAc sugar bound to the calcium binding site (PDB :6PY1). Calcium ions are shown in magenta.

MGL is a transmembrane protein composed of an intracellular signaling domain, a transmembrane domain, a coiled coil trimerization domain and a C-terminal Carbohydrate Recognition Domain (CRD) (Fig 1D). The CRD fold is highly conserved in C-type lectins and is organized as a double-loop structure (Fig 1E) stabilized by at least two conserved disulfide bridges. The overall domain is a huge loop in itself with its N- and C-terminus joined together, thanks to the first disulfide bridge, which contains another loop (the so-called long loop region) also stabilized by the second conserved cystine bridge (Zelensky and Gready, 2005). Some C-type lectin domains, including MGL, possess an additional N-terminal β-hairpin that is stabilized by a third cystine bridge conserved in these long-form sub-type of CRDs. The domain presents a mixed α/β fold and a large proportion of loops with undefined secondary structures (Fig 1E). For most of the CLRs reported, glycan binding site is calcium dependent and characterized by a tripeptide motif (EPN/QPD) and residues from the adjacent β-strand that assume metal coordination (Valverde et al., 2020). MGL possesses a QPD (267-269) motif characteristic of recognition of glycans with terminal galactoses (Fig 1F). The X-ray structure of human MGL-CRD (Gabba et al., 2021) in complex with galactose-containing ligands shows two galactose ring hydroxyl groups 3 and 4 bound to the calcium ion. Additionally, H286 is proposed to be responsible for selectivity towards N-Acetyl through a water mediated hydrogen bond (Marcelo et al., 2019). MGL binds preferentially to terminal N-acetyl Galactosamine residue and presents, for a C-type lectin, an unusually low (μM) dissociation constant for the monosaccharide (Diniz et al., 2019). The interaction of MGL with terminal galactoses from the core oligosaccharides was shown for *C. jejuni* LPS (van Sorge et al., 2009) and for *E. coli* R1 type core oligosaccharide (Fig 1C) (Maalej et al., 2019).

In this work, we have investigated MGL binding to oligosaccharides isolated from deacylated LPS or to native LPS directly exposed on whole cells. Our results show that in the trimeric oligomerized form, the carbohydrate recognition domain of MGL adopts a specific three-dimensional arrangement that allows a unique presentation of its 6 glycan binding sites (2 per CRD), composed of the canonical QPD calcium-binding motif and a newly described interaction site.

## RESULTS

### MGL Extracellular domain strongly binds to bacterial surface, independently of the QPD motif

MGL Extra-Cellular Domain (ECD) was shown by Nuclear Magnetic Resonance (NMR) to interact with the terminal galactoses of *E. coli* R1 type core OS. To establish MGL binding in the context of R1 oligosaccharide assembled at the cell surface, interaction of MGL-ECD was tested with live bacteria. *E. coli* bacteria exhibit variable structures of the core OS, so we chose to compare R1 and R3 types (Fig 1C, Fig S1) because they represent together more than 80% of *E. coli* strains including enterohemorrhagic species (Amor et al., 2000). Two bacterial strains carrying R1 and R3 core oligosaccharide structures but no O-antigen, respectively F470 and F653, were thus compared for MGL interaction. MGL-ECD was labelled with AlexaFluor 647 (AF647), incubated with *E. coli* bacteria and excess protein was washed. Bacteria were imaged by fluorescence microscopy. F470 bacteria were significantly labelled at their surface by MGL while F653 showed no labelling, confirming that MGL can recognize R1 core oligosaccharide on cells (Fig 2A). In order to ascertain that the interaction with the LPS observed was specific, the interaction was reproduced in presence of 10 mM GalNAc, that possess a low μM affinity for MGL, as a competitor and quantitatively assessed MGL binding by flow cytometry (Fig 2B and Fig S2). We found that GalNAc at high concentration could not significantly compete to the binding of MGL to R1 presenting cells.

**Figure 2.**
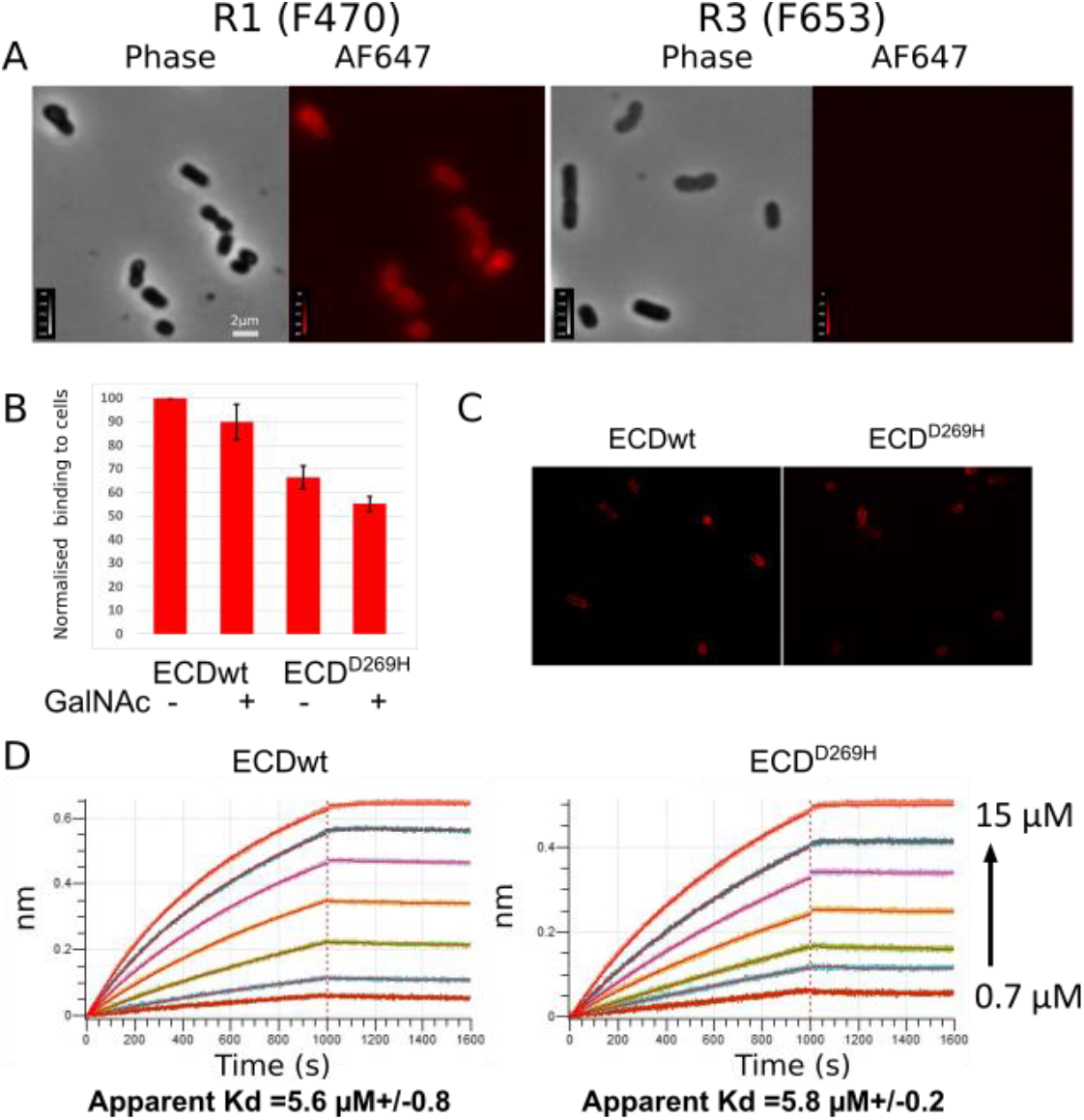
MGL ECD binds specifically to R1 presenting *E. coli* cells independently of the QPD motif. A) Phase contrast and epifluorescence microscopy images of AF647 labelled ECD incubated with R1 (left) or R3 (right) presenting *E. coli* cells. B) Flow cytometry quantification of MGL ECD wt and D269H variant labelled with AF647 bound to R1 cells in presence or absence of 10 mM GalNAc competitor. C) Confocal fluorescence image corresponding to conditions in B, showing strong MGL ECD^D269H^ binding to cells. D) *In vitro* interaction of immobilized MGL ECDwt and D269H variant to purified R1 LOS vesicles by biolayer interferometry.

The inability of the GalNAc monosaccharide to compete with the MGL binding to F470 cells could be ascribed to the multivalency of the interaction between the MGL trimer and R1 oligosaccharides presented on the cell surface. We thus designed a mutant of a key residue of MGL-CRD, that would abolish MGL carbohydrate binding capacity. D269, part of the conserved QPD motif (Fig 1F), is involved in calcium mediated binding of GalNAc to MGL (Diniz et al., 2019), so we decided to produce a D269H mutant to have a steric and electrostatic inhibition of the interaction with Ca^2+^ ion in canonical carbohydrate-binding site. MGL-ECD^D269H^ labelled with AF647 was thus incubated with F470 cells and imaged (Fig 2C). We surprisingly found that MGL^D269H^ was still able to significantly bind bacteria. This was quantified by flow cytometry that showed only a 30% decrease in binding of the D269H variant to cells (Fig 2B Fig S2) with little additive effect upon addition of 10 mM GalNAc. The behavior of this variant and the inability of GalNAc to inhibit significantly the binding suggest that, while the QPD motif is contributing to the interaction with R1 at the cell surface, it is not the main determinant of the interaction.

To exclude the interaction of MGL with another component at the cell surface of *E. coli* F470, the interaction between MGL and purified R1 LipoOligoSaccharide (LOS) has been tested *in vitro* by BioLayer Interferometry (BLI). MGL-ECD and MGL-ECD^D269H^ were biotinylated and immobilized on streptavidin coated BLI tips. R1 LOS was extracted and purified from F470 cells by Petroleum-Chloroform-Phenol (PCP) method and formed large vesicles in aqueous solution (de Castro et al., 2010). Injection of increasing concentrations of R1 LOS over immobilized MGL-ECD and MGL-ECD^D269H^, showed a significant interaction characterized by a slow association and a very slow dissociation. An apparent affinity constant in the low μM (around 5 μM) could be determined by steady-state analysis at the end of the association phase (Fig2 D Fig S3) and showed no affinity decrease for MGL-ECD^D269H^. The only difference observed with D269H variant was the maximum level of binding, Rmax, which was reduced by about 30%, similarly as observed on cells.

The *in vitro* interaction between purified partners confirmed that MGL was strongly binding to R1 core oligosaccharide on cells and *in vitro* when assembled as vesicles. While the integrity of the QPD motif contributes to the interaction with R1 core oligosaccharide, it is not the main determinant of the interaction. We thus hypothesized the existence of a secondary glycan binding site in MGL and investigated its localization by NMR.

### MGL carbohydrate recognition domain binds to LPS-derived oligosaccharides through a new binding surface

MGL-CRD and its binding to GalNAc and tumor-associated glycopeptides was previously characterized by NMR, X-ray crystallography and molecular dynamics. Those studies show a clear involvement of the QPD motif, with a particular contribution of H286 in the recognition of the N-acetyl moiety (Diniz et al., 2019; Gabba et al., 2021). MGL-CRD^wt^ and MGL-CRD^D269H^ have been produced and analyzed by ^1^H-^15^N NMR spectroscopy to localize the binding site of LPS-derived oligosaccharide. Wild-type MGL-CRD shows a spectrum similar to the one already published. D269H variant ^1^H-^15^N correlation spectrum is also characteristic of a well-folded protein and comparable to the wild-type spectrum (Fig S4). Backbone resonances of wild-type and D269H variant were assigned and used to predict their secondary structure content. It confirmed that MGL-CRD^D269H^ contains the same secondary structure elements than the wild-type protein (Fig S4). The mutation, by abolishing the proper coordination of the calcium ion probably destabilizes the whole GalNAc binding site. Therefore, the resonances from residues 265 to 282 remained unassigned in D269H variant.

First, the binding to GalNAc sugar was assessed for both proteins. 2D ^1^H-^15^N correlation experiments show resonances, each one corresponding to the amide frequencies of individual aminoacids. Addition of a ligand perturbs the amide frequencies at the vicinity of the binding site and can be good reporters of both the affinity and the aminoacids involved in the binding. As reported by Dinitz et al., MGL-CRD binds strongly to GalNAc in the characterized binding site between residues 264 and 296, with strong Chemical Shift Perturbations (CSP) of D269 and H286 amide resonances (Fig 3A Fig S5). MGL-CRD^D269H^, as predicted, does not show any CSP upon binding to GalNAc (Fig S6), consistent with its inability to bind to GalNAc affinity column during purification.

**Figure 3:**
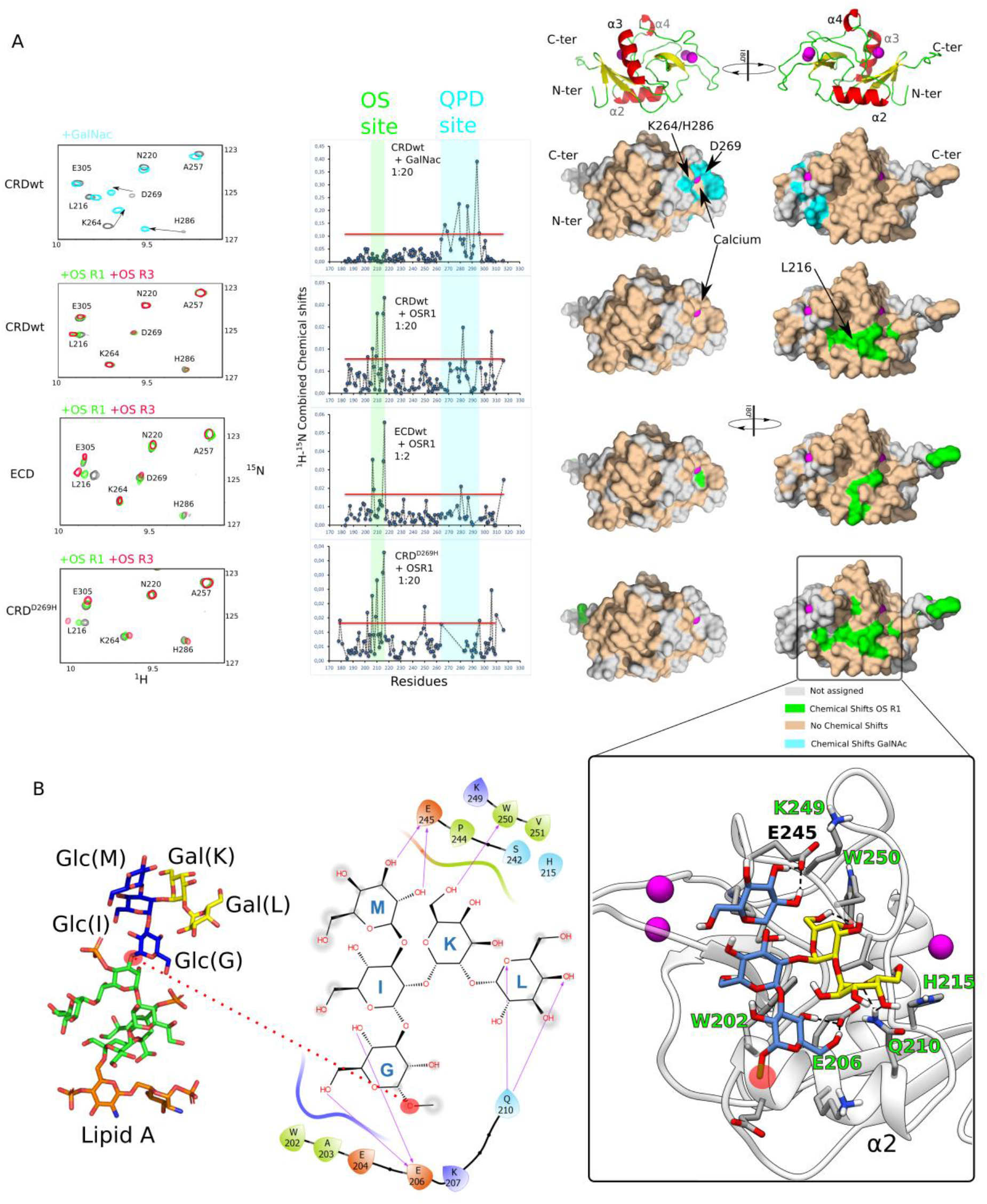
GalNAc and LPS derived oligosaccharides interact on two opposite surfaces of MGL. A) left: extracts of ^1^H-^15^N correlation spectra of the CRD, CRD^D269H^ and the CRD in the full ECD upon interaction with GalNAc, OS R1 or OS R3. Middle: CSP of the corresponding interactions represented with respect to aminoacid sequence. The red line marks the threshold of significant CSP. Right: Significant CSP represented on the CRD surface upon interaction with GalNAc or OS R1. B) Docking and molecular dynamics of R1 outer core pentasaccharide binding on the OS binding site. Left: Structure of OS R1. Middle: Two-dimensional schematic plot of the interactions between MGL and *E. coli* outer core pentasaccharide: solid arrows represent hydrogen bonds. Right: 3D representation of the MGL pentasaccharide complex. Residues of MGL involved in CSPs are labelled in green. Calcium ions are in magenta.

LipoOligoSaccharides assemble into large vesicles in solution that rapidly sediment and are not suitable to perform interactions by NMR. Soluble LOS derived oligosaccharides (OS) of R1 and R3 types (Fig 3B Fig S1) were then produced by chemical deacylation of LOS (Maalej et al., 2019). The interaction of CRD^wt^ and CRD^D269H^ was then tested with R1 and R3 OSs (Fig 3 Fig S5,6,9). Interaction with OS R1 showed CSP of the CRD^wt 1^H-^15^N resonances on a fast exchange regime with respect to NMR timescale with no saturation of the binding even at high OS concentration, suggesting a weak affinity (Kd ≥ 5 mM). Furthermore, residues of the CRD experiencing high CSP upon OS R1 binding lie on a surface opposite to the GalNAc binding site, in green on figure 3A, and involve residues 202-216 around the α2 helix. The same interaction performed with the D269H variant showed a very similar interaction site opposite from the QPD motif. We thus postulate that the new interface perturbed by OS R1 is responsible for the binding of MGL to F470 *E. coli* and to purified R1 LOS vesicles. As a control, we also tested the binding of the CRD wt and D269H variant to OS R3 in the same conditions (Fig 3 and Fig S9). OS R3 caused very similar CSP at the surface of CRD^wt^ and CRD^D269H^. The new interaction surface of MGL involved in glycan binding does not show specificity for R1 core oligosaccharide on the contrary to results obtained on cells. The configuration of the NMR interactions is very different from the *in vivo* experiments; the CRD domain is used instead of the ECD and the OSs are free in solution and are not presented on the cell surface as multivalent ligands. In order to confirm that the binding observed on the isolated CRD also applies to the CRD in the context of the trimeric ECD, we investigated the ECD by NMR.

The extracellular domain of MGL is a large protein (homo-trimer of 84kDa) for NMR spectroscopy due to signal broadening arising for long molecular tumbling correlation times. The protein was thus expressed and purified as a perdeuterated version. This is to our knowledge the first example of a full-length C-type lectin ECD studied by NMR. ^1^H-^15^N correlation spectrum of ^2^H,^15^N-labeled MGL-ECD is of high quality considering the protein size and elongated shape and is characteristic of a well-folded protein. When comparing the ^1^H-^15^N resonances observed on spectra recorded with isolated CRD, it is apparent that the footprint of the CRD domain is present in the ECD of MGL (Fig S7). Several additional overlapped resonances can be observed around 8.2 ppm in the proton dimension and probably arise from the coiled-coil domain. The low stability (several days) of MGL-ECD at the temperature needed to record NMR spectra (above 35°C) did not allow its *de novo* assignment by backbone assignment experiments. The good ^1^H and ^15^N agreement between CRD signals in ECD and isolated CRD permitted the transfer of most assignments from CRD to ECD (Fig S7). MGL-ECD was thus titrated with increasing concentrations of OS R1 and OS R3. CSP induced by the interaction with R1 or R3 OS are very similar to those observed with isolated CRD, though the surface involved is not as extended (Fig 3A, Fig S8). This suggests that the assembly of the CRD domain in the full length ECD has no influence on the selectivity of MGL towards either R1 or R3 chemical structure when interaction occurs with isolated oligosaccharides.

Previous report showed that the outer core galactose sugars of R1 oligosaccharide were selectively recognized by MGL (Maalej et al., 2019). The outer core pentasaccharide of R1-OS was thus docked (See methods section) onto the glycan site shown in that study, followed by a molecular dynamics simulation (Fig 3 B and Fig S10). The pentasaccharide stayed on the binding site for more than half of the simulation time. E245 and E206 interact with 2- and 3-OH, and 4- and 6-OH moieties of glucose M and G respectively. Additionally, Q210 is in charge of the interaction with hydroxyl moiety at position 4 of galactose L, whereas galactose residue K interacts with W250 (Fig 3C). The pose of the ligand is compatible with the NMR interaction experiments performed here and the residues experiencing CSP in NMR titrations, in particular Q210 and E206. Interestingly, and in accordance with the low affinity of OS R1 for MGL, at the end of the MD simulation the pentasaccharide did not longer remain on the secondary binding site and started to shift towards the GalNAc binding site.

The presence of two different glycan binding sites at the surface of the MGL-CRD on two opposite surfaces is unprecedented in C-type lectins. It suggests that in the extracellular domain both sites are accessible to bind their ligands, we thus investigated the global arrangement of the CRDs in MGL-ECD.

### MGL-CRDs are oriented perpendicular to the coiled coil domain and can present six sugar binding sites to bacterial surfaces

The structure of MGL-ECD is unknown and we studied its overall structure by Small Angle X-ray Scattering. This method enables to assess the size and shape of a macromolecule in solution, at a low resolution. MGL-ECD SAXS scattering curve confirms the presence of a trimeric protein with an estimated MW of 94 kDa (vs 84 kDa Theoretical MW), and a gyration radius of 5.6 nm, suggesting an elongated protein(Smilgies and Folta-Stogniew, 2015). Calculation of Pairwise distribution P(r) showed a maximum interatomic distance of 17 nm (Fig S11) consistent with the expected elongated shape of the ECD. P(r) was used to calculate an envelope of MGL-ECD (See Methods section). The envelope (Fig 4A) is characterized by an elongated structure, corresponding to the coiled-coil domain, with three large bulges on its side that can be ascribed to the CRDs. The SAXS-derived envelope does not allow to orient at an atomic scale the CRD, but the location of the bulges suggests that the CRD domains are perpendicular to the coiled-coil domain. This orientation would be significantly different from an about 120° angle observed between coiled-coil and CRD domain observed for langerin or MBP trimers (Feinberg et al., 2010; Ng et al., 2002).

**Figure 4.**
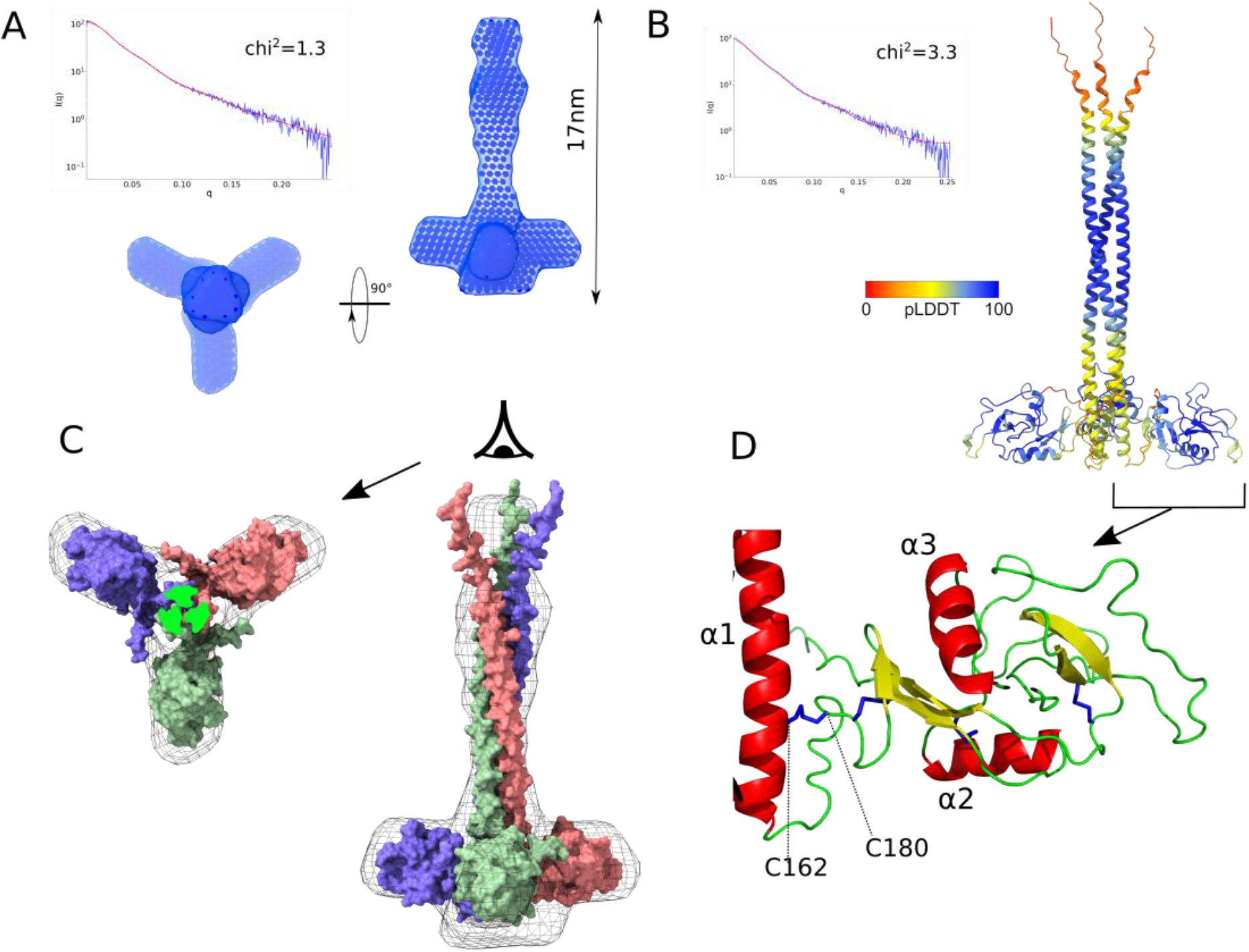
Combined SAX-alphafold model of MGL-ECD A) SAXS of MGL-ECD with SAXS curve (top left blue) and the corresponding fit (red) of the SAXS envelope, calculated from P(r) distribution, shown as surface from side and N-terminus of coiled-coil view. B) Alphafold model with the best correspondence to SAXS curve, colored by pLDDT score. The calculated SAXS curve of this model is shown (red) compared to the experimental curve (blue). C) Best alphafold model of MGL-ECD adjusted into the SAXS enveloppe (in mesh) in side and from the N-terminus of coiled-coil view. D) Closeup view on the C162-C180 disulfide bond orienting the CRD in the best alphafold model.

To position the CRD into the SAXS envelope, MGL-ECD models were generated with the alphafold structure prediction protocol (Jumper et al., 2021; Tunyasuvunakool et al., 2021) (see methods section). This method has provided atomic scale prediction of protein structure of unprecedented accuracy with a combination of machine-learning and evolutionary data. The models show a long N-terminal coiled-coil domain (N86-N169) followed by the CRD (C181-H316). The arrangement of the CRD relative to the coiled-coil domain is variable and allows to sort the models into two clusters. The lack of well-defined interdomain contacts can be explained by low Alphafold per residue score (pLDDT) at the interface and little interactions predicted in the Prediction Alignment Error matrix (Fig S12). One new disulfide bond is nevertheless predicted in all models between coiled-coil (C162) and CRD (C180) (Fig 4D Fig S12). This disulfide bond is consistent with Mass Spectrometry analysis of MGL-ECD which displays an 8 Da difference with the theoretical mass, corresponding to a total of 4 disulfide bonds (Fig S13). The two cysteines involved are also strictly conserved in the MGL family in mammals (Fig S14 Table S1) and the disulfide bond at the corresponding position was shown experimentally in the homologous protein asialoglycoprotein receptor 1 (Ruiz and Drickamer, 1996).

The two clusters of models are different in the orientation of the CRD domains with an almost 180° rotation around the C160-C182 disulfide bond (Fig S12). To determine which cluster corresponds better to the conformation in solution, models were evaluated against experimental SAXS data. SAXS curves were back-calculated from the models and compared to the experimental one (Fig 4B Table S2 and Fig S15). Cluster 2 structures show systematically a better fit compared to cluster 1. The best matching structures of each cluster can also be inspected visually by adjusting the structures into the SAXS-derived envelope (Fig 4C, Fig S16). The cluster 2 models are in the best accord to the SAXS data and the structure with the lowest chi^2^ with respect to the SAXS curve was retained for analysis (Fig 4 B-D).

The two glycan binding sites of the CRD, the canonical QPD motif and the newly described OS binding site, can be represented on the surface of the MGL model (Fig 5A) in cyan and green, respectively. The orientation of the CRD is such that QPD and OS sites from two neighboring CRDs face each other. If we consider that the most likely configuration of MGL binding to the bacterial surface would be perpendicular to the membrane, the CRDs are able to present up to six glycan binding sites to LPS core oligosaccharides (Fig 5B). In that configuration, even if the affinity of MGL for isolated core OS is low, the avidity of the interaction would ensure a tight binding to the surface, consistent with our observations on bacteria presenting R1 core oligosaccharides.

**Figure 5.**
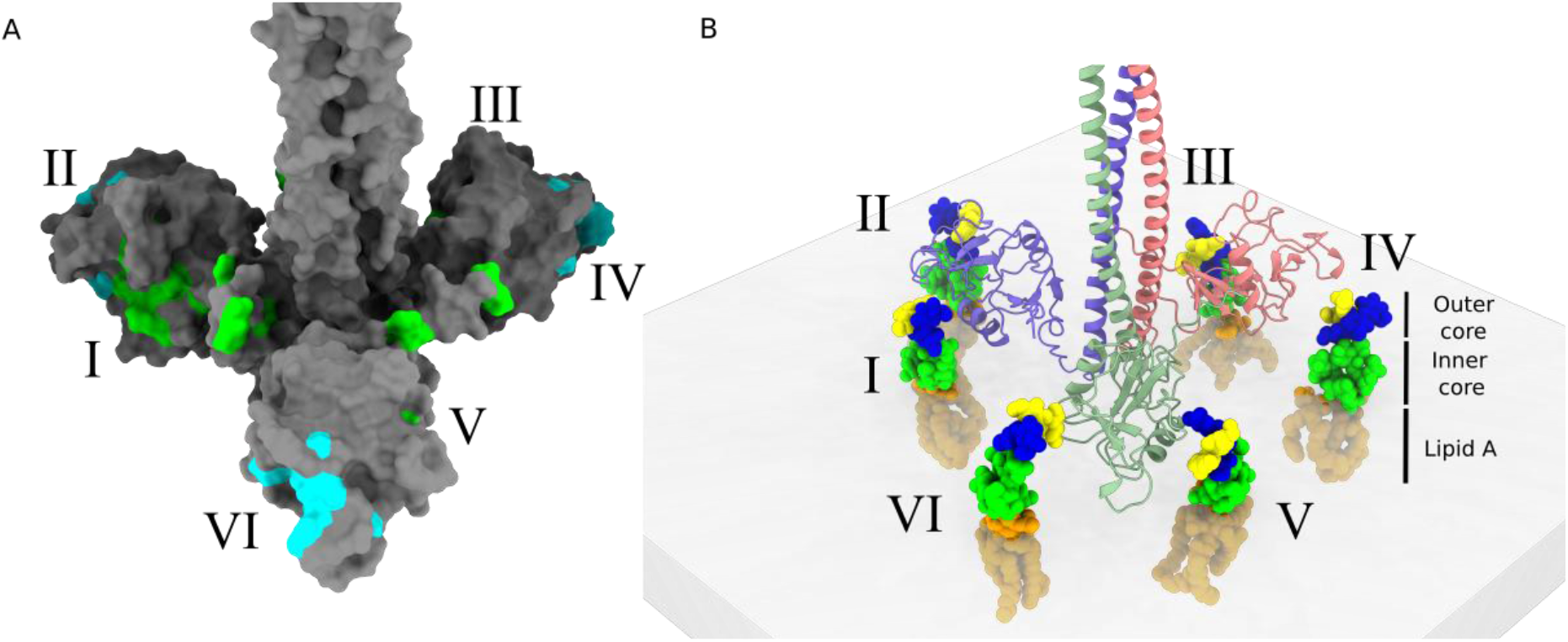
Combination of SAXS and alphafold defines a CRD arrangement that presents up to six accessible glycan binding sites (I to VI). A) Representation of the two glycan binding sites, with the NMR CSP of GalNAc(cyan) and OS R1(green) on the best SAXS-alphafold MGL structure. B) Schematic view of 6 R1 LOS molecules, with an orientation similar to what is found at the bacterial surface, facing the 6 glycan binding sites of MGL-ECD.

## DISCUSSION

Glycoconjugates are present at the surface of most cells, as well as in extracellular matrices and biofilms. In complex multicellular organisms the sugar environment is very rich and heterogeneous. The immune system must recognize friends from foes and clear pathogens, but also tailor its response to avoid excessive inflammatory response. The recognition of pathogens vs commensals is critical and also relies on subtle variations of microbial glycome. MGL has been reported so far to recognize several bacterial pathogens, with different cell wall structures, through their surface glycans.

While MGL attaches strongly to *E. coli* surface presenting R1 type core oligosaccharides, this binding is largely independent of the QPD GalNAc binding site. We could show that a second interface, opposite to the QPD binding site, binds LPS core oligosaccharides. Several examples exist of secondary binding sites in C-type lectins. They can be located adjacent to the conserved calcium binding site to extend the binding interface and confer specificity towards a given ligand like observed for Trehalose Di-mycolates for Mincle (Furukawa et al., 2013), or on a more remote site like for Heparin for Langerin (Chabrol et al., 2012) and through cooperativity for DC-SIGN (Medve et al., 2019; Wawrzinek et al., 2021). The presence of a second binding site completely opposite to the canonical binding site is nevertheless unusual. We suggest that this is correlated with the peculiar 3D arrangement of MGL-CRDs compared to other multimeric C-type lectins. Other trimeric C-type lectins like Langerin or Mannose-Binding Protein (MBP) (Feinberg et al., 2010; Ng et al., 2002) adopt a compact arrangement of their CRDs (Fig 6) with their canonical binding sites accessible at the extremity of the proteins. Their calcium binding sites lie within 50 Å of each other compared to about 80 Å for MGL. It allows MGL to target surfaces with much distant glycan epitope. Furthermore, this extended conformation makes the C-terminal loop of the coiled-coil neck domain, connecting it to the CRD, accessible at the surface and could contribute to glycans binding (Fig 6). This region of the protein varies between isoforms1 and 2 of human MGL (Fig 6 Fig S14) with insertion of 3 additional residues (G171-E172-E173) in isoform 2 (this study). These residues could participate to the interaction of MGL with a bacterial surface but could also be important for the orientation of the CRD. The reduction of the coiled-coil CRD linker in isoform 1 would alter, in turn, the orientation of the CRDs by likely leading them to rise upward. Thus, while this 3 residues insertion, from the isoform 1 to 2 of MGL, do not modify the glycan binding specificities of their CRDs, it might impact drastically the relative geometry of the CRDs in both trimeric isoform and thus their specificity towards different glycan landscape. The conserved disulfide bond positions the CRD domain perpendicular to the coiled-coil axis and has important implications with respect to glycan binding. As we have recently shown on another CLR, thanks to molecular dynamic studies, DC-SIGN can adapt to various distance distribution of glycan epitope presentation thanks to a rather large flexibility between the neck and the CRDs domains (Porkolab et al., 2023). Here a different situation occurs in the case of MGL. The presence of the newly identified C162-C180 bridge, strongly constrains the extension capabilities of CRDs from the neck (Fig 4D). However, the CRD domains show here no extensive contacts with the coiled-coiled neck domain and subtle variations of the CRD orientation through rotation around the disulfide bonds axis might allow plasticity in the presentation of the binding sites. However, the limitation in distance is compensated here, in MGL, by the presence of the additional non canonical OS binding site on the opposite side, within the CRD, of the Ca^2+^-dependent QPD site. This, combined to CRDs subtle rotation, might provide a large set of potential adaptation to different surfaces.

**Figure 6.**
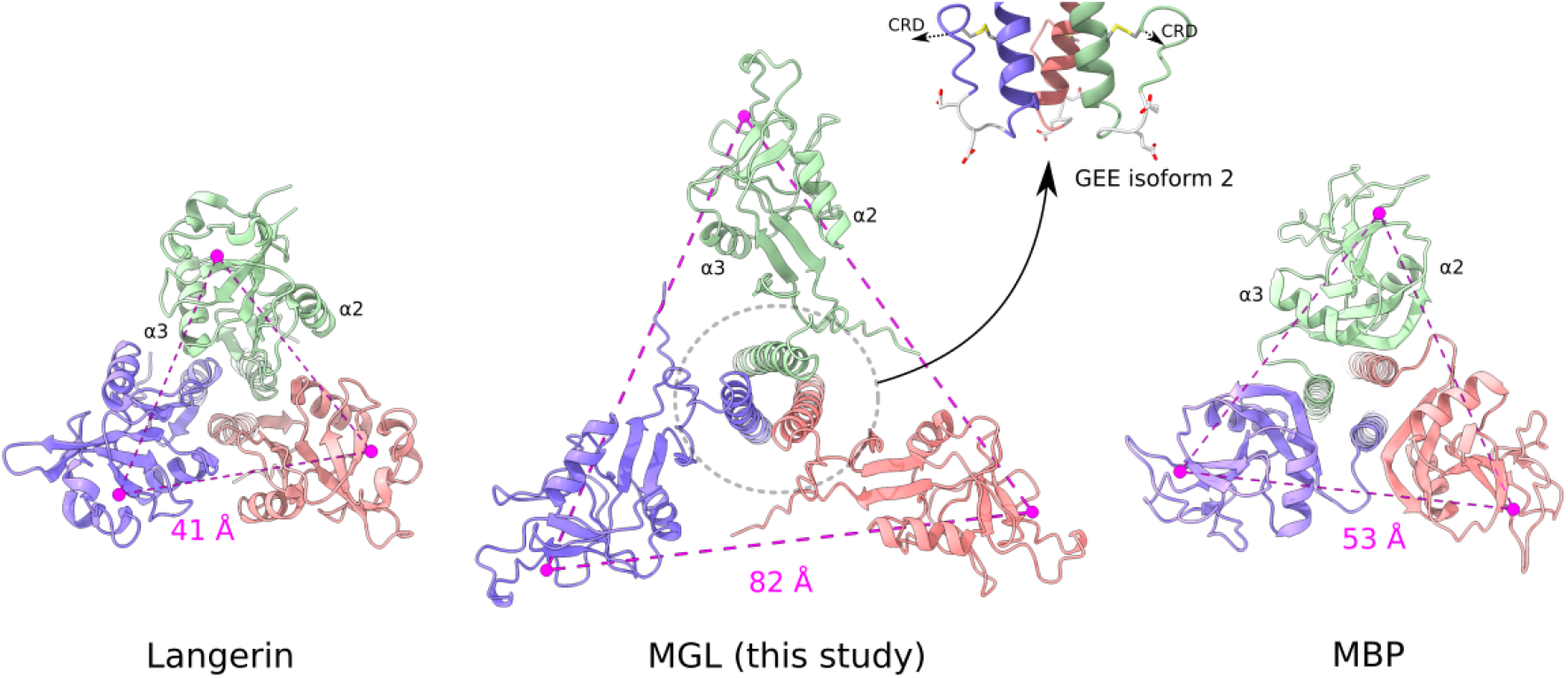
Comparison of the MGL model with other trimeric C-type lectins. The MGL model, Langerin (PDB:3KQG) and Mannose Binding Protein A (PDB:1KWW) are shown from the C-terminal side of the coiled-coil domain. The calcium ions of the canonical binding sites are indicated as well as the distances in magenta between adjacent sites. The C-terminus of the coiled-coil domain of MGL is surface accessible (grey circle) and the MGL isoform 2 studied here has an additional GEE tripeptide (grey). The C162-C180 disulfide bond is in yellow.

Here, the CRDs orientation makes both QPD and OS binding sites accessible for binding glycans assembled on a surface. The presence of these six binding sites highly increases the multivalency of the interaction and probably explains the very broad Pathogen Associated Molecular Patterns that MGL is capable to recognize, from both gram positive and negative bacteria, as well as *M. tuberculosis* (Mnich et al., 2019; Naqvi et al., 2021; van Sorge et al., 2009; Vliet et al., 2009). We can hypothesize that the mode of recognition by MGL of Teichoic Acids, that are polysaccharides assembled similarly as LPS at the surface of *S. aureus* (Mnich et al., 2019), resembles that of LPS. So far, we have examined the binding of MGL to LPS that do not contain O-antigens. Most clinically relevant gram-negative bacteria possess O-antigen of very variable compositions and length (Whitfield et al., 2020). This dense and long (∼10-40nm) layer of polysaccharides could be either recognized by MGL, thanks to its ability to bind various glycans, or on the other hand it could block access to the core oligosaccharides and prevent recognition. This should be the focus of future studies on the role of MGL in the recognition of gram-negative bacteria and the subsequent implications in the regulation of the immune response.

## Material and methods

### Protein expression and purification

Human MGL isoform 2 ECD (residues Q85-H316) with an N-terminal StrepTagII and a Factor Xa cleavage site was expressed and purified as already reported (Maalej et al., 2019). Briefly, MGL-ECD was over-expressed in *E. coli* BL21(DE3) cells in inclusion bodies. Inclusion bodies were solubilized in guanidine buffer (25 mM TRIS pH 8, 150 mM NaCl, 6 M Guanidine, 0.01% B-mercaptoethanol). MGL-ECD was subsequently refolded using a drop-by-drop dilution in renaturation buffer (100 mM TRIS pH 8, 150 mM NaCl, 25 mM CaCl_2_) and was subjected to 2 purification steps; a GalNAc-Agarose affinity column (Sigma), eluted with EDTA buffer (150 mM NaCl, 25 mm TRIS pH 8, 10 mM EDTA) followed by a Toyopearl HW-50S gel filtration column (Tosoh Bioscience). MGL-ECD was also produced as perdeuterated ^2^H, ^15^N-labelled form ([U-^2^H,^15^N] MGL-ECD) in 95% D2O with D-glucose-d7 as glucose source as described (Jean et al., 2014). MGL-ECD^D269H^ mutant was over-expressed in *E. coli* BL21(DE3) cells in LB medium as inclusion bodies which were subjected to the same solubilization, and renaturation steps described above. MGL-ECD^D269H^ was purified using an AktaXpress with a Strep tag affinity column eluted with 2.5 mM Desthiobiotin followed by a Toyopearl HW-50S gel filtration column (Tosoh Bioscience).

MGL-CRD and MGL-CRD^D269H^ (C181-H316) with N-terminal His-Tag and TEV cleavage site were expressed and purified as described (Bulteau et al., 2022) in M9 minimal medium as ^13^C,^15^N labelled proteins. An MGL-CRD^D269H^ binding assay was performed on the GalNAc-Agarose affinity column used for ECD purification to assess its affinity for GalNAc, which revealed it did not bind to GalNAc affinity column.

### Fluorescence microscopy, flow cytometry

MGL-ECD and MGL-ECD^D269H^ were labelled with Alexafluor647-NHS (Invitrogen). Briefly MGL at 5 mg/ml in PBS buffer was incubated in 200 mM sodium bicarbonate and 0.4 mg/ml AF647-NHS for one hour. Excess dye was removed with G25-PD10 desalting column (GE Healthcare) and MGL fractions dialyzed further against PBS buffer and concentrated. *E. coli* R1 bacteria carrying R1 core oligosaccharide (F470 derivative from *E. coli* O8:K27), and R3 (F653, derivative from *E. coli* O14:K7) (Amor et al., 2000) were grown in LB at 37°C under agitation up to 0.9 OD_600nm_. Cells were collected by centrifugation, washed in cold PBS and incubated with 670 nM MGL-AF647 in PBS, 2 mM CaCl_2_ buffer for 15 min. Cells were washed five times with cold PBS and imaged. For each sample, 2 μL of cells in suspension were mounted between a glass slide and a 1.5H 170μm thick glass coverslip, and observed using an inverted IX83 microscope, with the UPLFLN 100× oil immersion objective from Olympus (numerical aperture 1.49), using a fibered Xcite™ Metal-Halide excitation lamp in conjunction with the appropriate excitation filters, dichroic mirrors, and emission filters specific for AF647 (4×4MB set, Semrock). Acquisitions were performed with Volocity software (Quorum Technologies) using a sCMOS 2048 × 2048 camera (Hamamatsu ORCA Flash 4, 16 bits/pixel) achieving a final magnification of 64 nm per pixel.

Flow cytometry was performed on a VYB device (Miltenyi biotech) and analyzed with Macsquant software. Cells (50μl) grown in LB at DO_600nm_=1 were resuspended in presence of 670 nM MGL-AF647 (wt or D269H variant) in PBS, 2 mM CaCl_2_ with/without 10 mM GalNAc for 15 min, centrifuged twice to remove excess protein, resuspended in 150 μl and injected for FACS analysis until 200000 events were recorded. MGL ECD binding to cells was expressed as % population x mean fluorescence (cy5 channel) and normalized to 100 % for MGL ECD wt binding.

### LOS and OS preparation

F470 and F653 cells were grown in LB. LipoOligosaccharides were extracted following the PCP method and de N- and O-acylated as already described (de Castro et al., 2010; Maalej et al., 2019).

### Bio-Layer Interferometry (BLI) experiments

ECD^wt^ and ECD^D269H^ were biotinylated on carboxyls with EDC and Biotin Hydrazide. Protein at 2 mg/mL in interaction buffer (150 mM NaCl 50 mM Phosphate pH 8) was treated with 1.25 mM Biotin LC-Hydrazide (Thermofisher) and 5 mM EDC for two hours then dialyzed against the initial buffer. The interferometry measurements were carried out on an octet Red96(Fortébio) using streptavidine coated biosensors (SA). Proteins were immobilized at 5μg/ml in HEPES buffered saline buffer, 0.02 % Tween20 to reach 1 nm of response. The functionalized sensors were equilibrated in interaction buffer (50 mM phosphate 150 mM NaCl pH 8), then immersed in the wells containing R1 LOS vesicles at concentrations ranging from 0.7 to 15 μM with agitation for 1000 s (association) then immersed in buffer for 600 s (dissociation). The data obtained were processed with the octet evaluation software after subtraction of reference non-functionalized biosensors (SA) to remove the contribution of non-specific interactions.

### NMR titrations

Human ^15^N labelled MGL-CRD^wt^ or MGL-CRD^D269H^ at 50μM in 25 mM TRIS pH 8, 150 mM NaCl, 4 mm CaCl_2_ was titrated with increasing concentrations of GalNAc, R1 or R3 Oligosaccharides up to 20 molar equivalents glycan:CRD. ^1^H-^15^N-BEST-TROSY correlation experiments were recorded at 30°C on an 850, 700 or 600 MHz Bruker NMR spectrometer equipped with a cryoprobe at each oligosaccharide addition. NMR titration experiments with MGL-ECD were performed at a concentration of 600 μM of the ^2^H,^15^N MGL ECD with 1 and 2 molecular equivalents of either OS R1 or R3 ligands. ^1^H-^15^N-BEST-TROSY correlation spectra were collected at 35°C on Bruker Avance spectrometer at 850 MHz. All spectra were processed using TopSpin 3.5 software and analyzed using CcpNmr analysis 3.0 software. CSP, corresponding to the chemical shift change in the ^1^H–^15^N BTROSY spectra upon addition of ligands were calculated as CSP=((Δδ^1^H)^2^+([Δδ^15^N/10])^2^)^1/2^, where Δδ^1^H and Δδ^15^N are chemical shift changes in amide proton and amide nitrogen, respectively. CSPs higher than twice the standard deviation of all chemical shifts were considered significant.

### SAXS

SAXS data have been recorded on MGL-ECD domain at 1 mg/ml in 25 mM TRIS pH 8, 150 mM NaCl, 4 mm CaCl_2_ buffer at 25°C at European Synchrotron Radiation Facility (ESRF) BM29 Biosaxs beamline (Grenoble). Automatic frames selection and buffer subtraction was performed by ISPyB (de Maria Antolinos et al., 2015). SAXS data were analyzed with Atsas 3.1.3 (Manalastas-Cantos et al., 2021) and BIoXTAS RAW (Hopkins et al., 2017). P(r) distribution function was used as input for DAMMIF online, doing five runs including P3 symmetry and prolate anisotropy. The five solutions were sorted by DAMAVER as two clusters and the most representative envelope of the best cluster is presented. Alphafold multimer was run with the entire sequence of the MGL-ECD construct expressed, and as a trimeric protein as input. 24 models have been generated and the 10 best ranked models according to their DockQ score, were retained for further analysis (Basu and Wallner, 2016).

### Docking and Molecular Dynamics simulation

Docking calculations were performed using AutoDock 4.2.2 and analyzed with AutoDockTools (Morris et al., 2009). The ligand was downloaded from GLYCAM website (www.glycam.org) and all rotatable bonds were set as free to move during calculations. The grid point spacing was set to 0.375 Å, and a hexahedral box was built with x, y, z dimensions: 60 Å, 50 Å, 60 Å centered in the centroid position among the binding pocket of MGL residues. In total, 200 runs using Lamarckian Genetic algorithm were performed, with a population size of 150, and the maximum number of energy evaluations set at 2500000. After docking, the 200 poses were clustered in groups with root-mean-square deviation less than 2.0 Å and the clusters ranked according to the lowest energy representative of each cluster. Molecular dynamic calculations were performed with AMBER 18 (Case et al., 2018) software in explicit water by using AMBER ff14SB, Glycam06j-1 and TIP3P force fields for the protein residues, the saccharide ligand and the water solvent molecules respectively. In order to prepare the protein, missing hydrogen atoms were added, and protonation state of ionizable groups and cap termini were computed by using Maestro Protein Preparation Wizard (Schrödinger Release 2022-3, Maestro, Schrödinger, LLC: New York, 2021). The MGL-pentasaccharide system was then hydrated with an octahedral box containing the explicit TIP3P water molecules buffered at 10 Å, adding also counterions to neutralize the system. The input files were generated using the tleap modules of the AMBER package. The minimization steps were performed using Sander module while molecular dynamic calculations were performed using the PMEMD module. At this point, an energy minimization process was performed to refine the initial structure. The calculations employed SHAKE for the C-H bonds and 1 fs of integration step. Periodic boundary conditions were applied, as well as the smooth particle mesh Ewald method to represent the electrostatic interactions, with a grid space of 1 Å. The system was minimized, at first, holding the complex, while a second minimization was performed on the entire system. Furthermore, the whole system was slowly heated from 0 to 300 K applying a weak restrain on the solute. Temperature was increased from 0 K to 100 K at constant volume then, from 100 K to 300 K, in an isobaric ensemble. Thereafter, temperature was kept constant at 300 K during 50 ps with progressive energy minimizations and solute restraint. Once completed the equilibration, the system restraints were removed, and the systems then advanced in an isothermal-isobaric ensemble along the production. The system coordinates were saved and used for the 100 ns simulations using the PMEMD module implemented in AMBER. Coordinate trajectories were recorded each 2 ps throughout production runs, yielding an ensemble of 10000 structures. Trajectories were analyzed using the ptraj module within AMBER 18 and VMD (Humphrey et al., 1996) program was used to visualize the MD results. Each trajectory was submitted to cluster analysis with respect to the ligand RMSD using K-mean algorithm implemented in ptraj module. The representative structure of the most populated cluster was considered to depict the complex interactions.

## Acknowledgements

We would like to thank A. Imberty and N. Thielens for stimulating discussions. We thank the Agence Nationale de la Recherche (ANR) PIA for Glyco@Alps (ANR-15-IDEX-02) and their support of M.M. and C.L. We thank Rose-Laure Revel-Goyet, Françoise Lacroix, Oleksandr Glushonkov and Jean-Philippe Kleman (IBS, Grenoble) for the support and access to the Cell imaging Platform. This work used the platforms of the Grenoble Instruct-ERIC center (ISBG; UAR 3518 CNRS-CEA-UGA-EMBL) within the Grenoble Partnership for Structural Biology (PSB), supported by FRISBI (ANR-10-INBS-0005-02). M.A. received funding from GRAL, the Grenoble Alliance for Integrated Structural and Cell Biology, a program of the Chemistry Biology Health Graduate School of Université Grenoble Alpes (ANR-17-EURE-0003). This project has received funding from the European Research Council (ERC) under the European Union’s Horizon 2020 research and innovation program under grant agreement No 851356 to R.M. This work was granted access to the CCRT High-Performance Computing (HPC) facility under the Grant CCRT2022-lagurice awarded by the Fundamental Research Division (DRF) of CEA.

