## supplementary information for "The unique three-dimensional arrangement of Macrophage Galactose Lectin enables *E. coli* LipoPolySaccharides recognition through two distinct interfaces"

### Supplementary information for Abbas, Maalej et al.

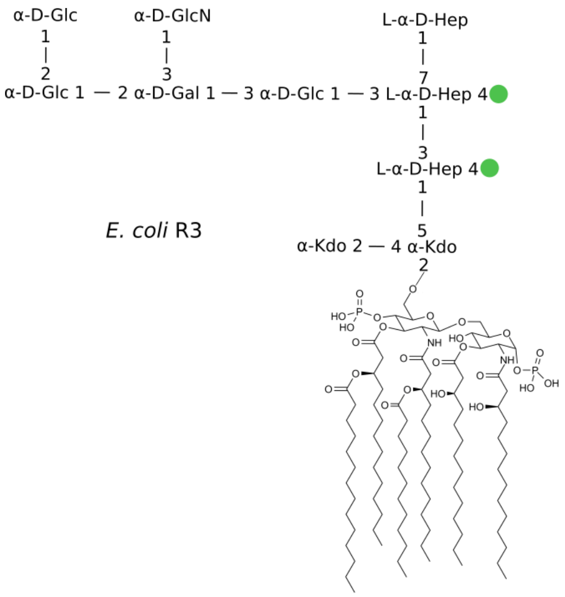


Figure S1: Structure of the main form of *E. coli* R3 type LipoOligoSaccharide

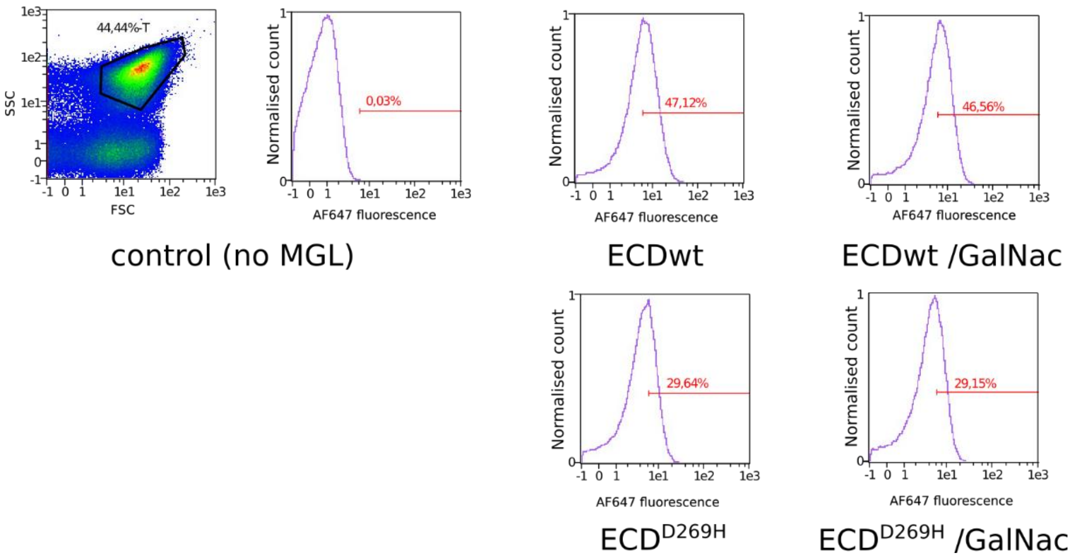


Figure S2 : Flow Cytometry of F470 cells with/without ECDwt and ECD$^{D269H}$ labelled with Alexafluor647. Left selection of bacterial population on 2D representation of Forward (FSC) and side (SSC) scattering and on the right count vs fluorescence of the selected bacterial population. Only the population within the red range was considered to have significant fluorescence. Associated % of population is indicated (before normalization to 100% for ECDwt shown in Fig 2).

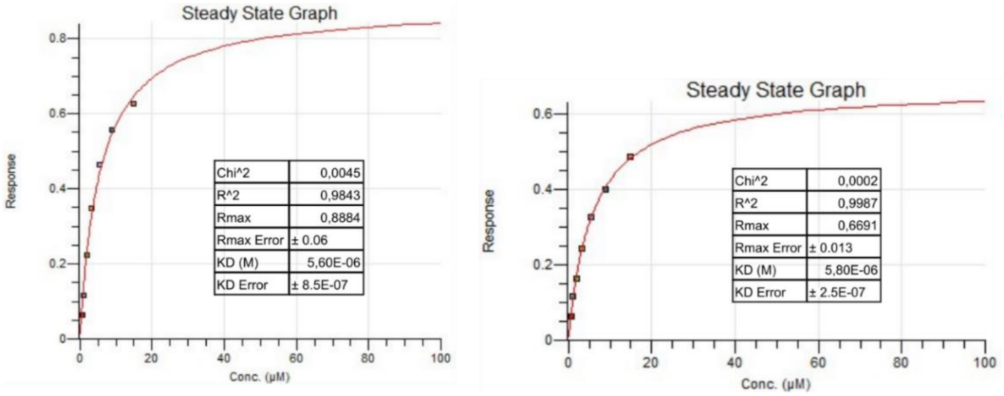


Figure S3 : Biolayer interferometry steady-state Kd determination for $ECD_{wt}$ (left) and $ECD^{D269H}$(right) with LOS R1 as ligand.

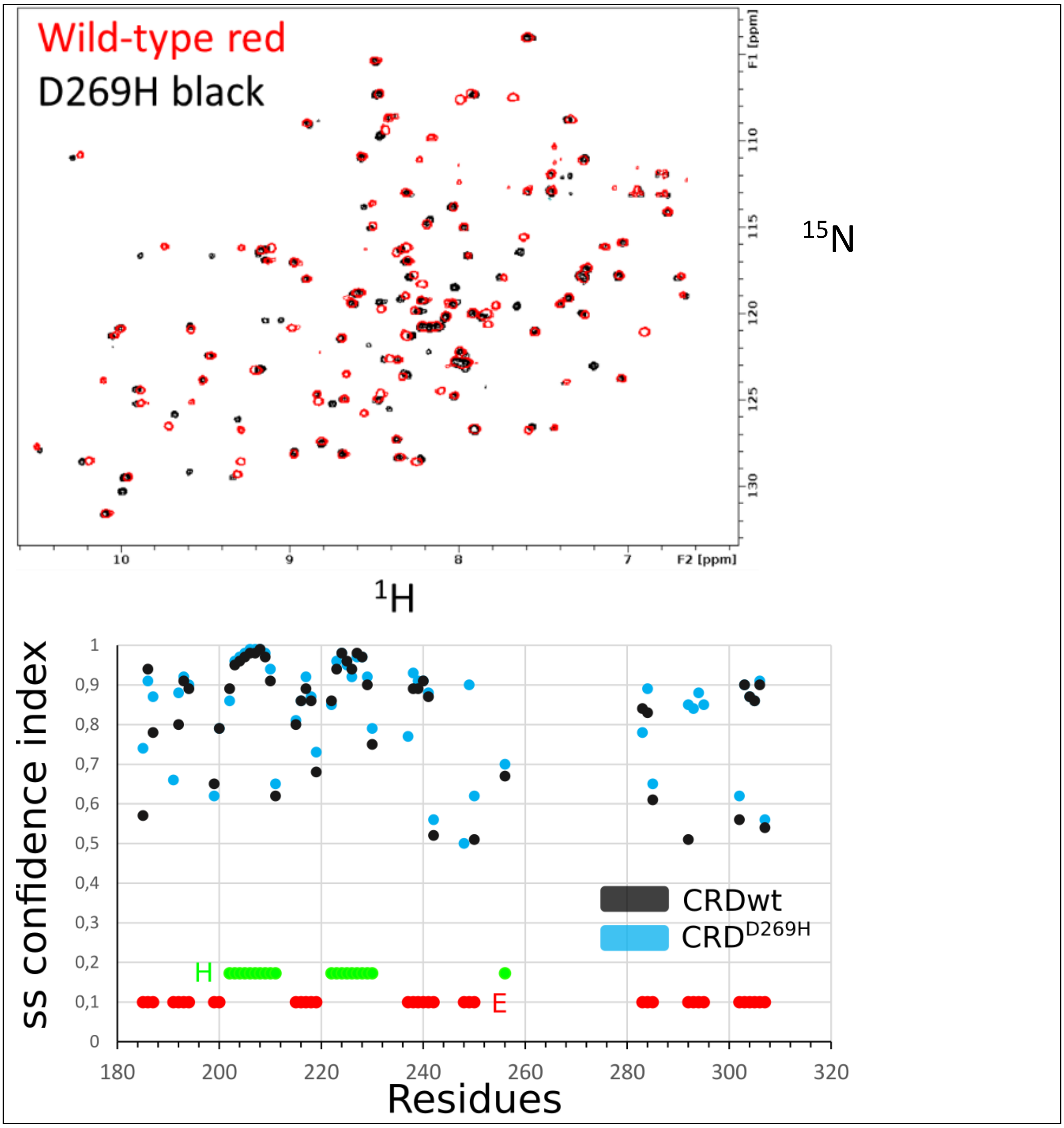


Figure S4 : Comparison of MGL-CRD wt and D269H variant by NMR. Top Overlay of Best-Trosy $^{15}$N-$^{1}$H correlation spectra of CRD$^{wt}$ and CRD$^{D269H}$ at 30°C. Bottom: Secondary structure prediction by Talos-n(Shen and Bax, 2013) from backbone chemical shifts of CRD and the D269H variant. Helix(H) or extended(E) predictions are shown.

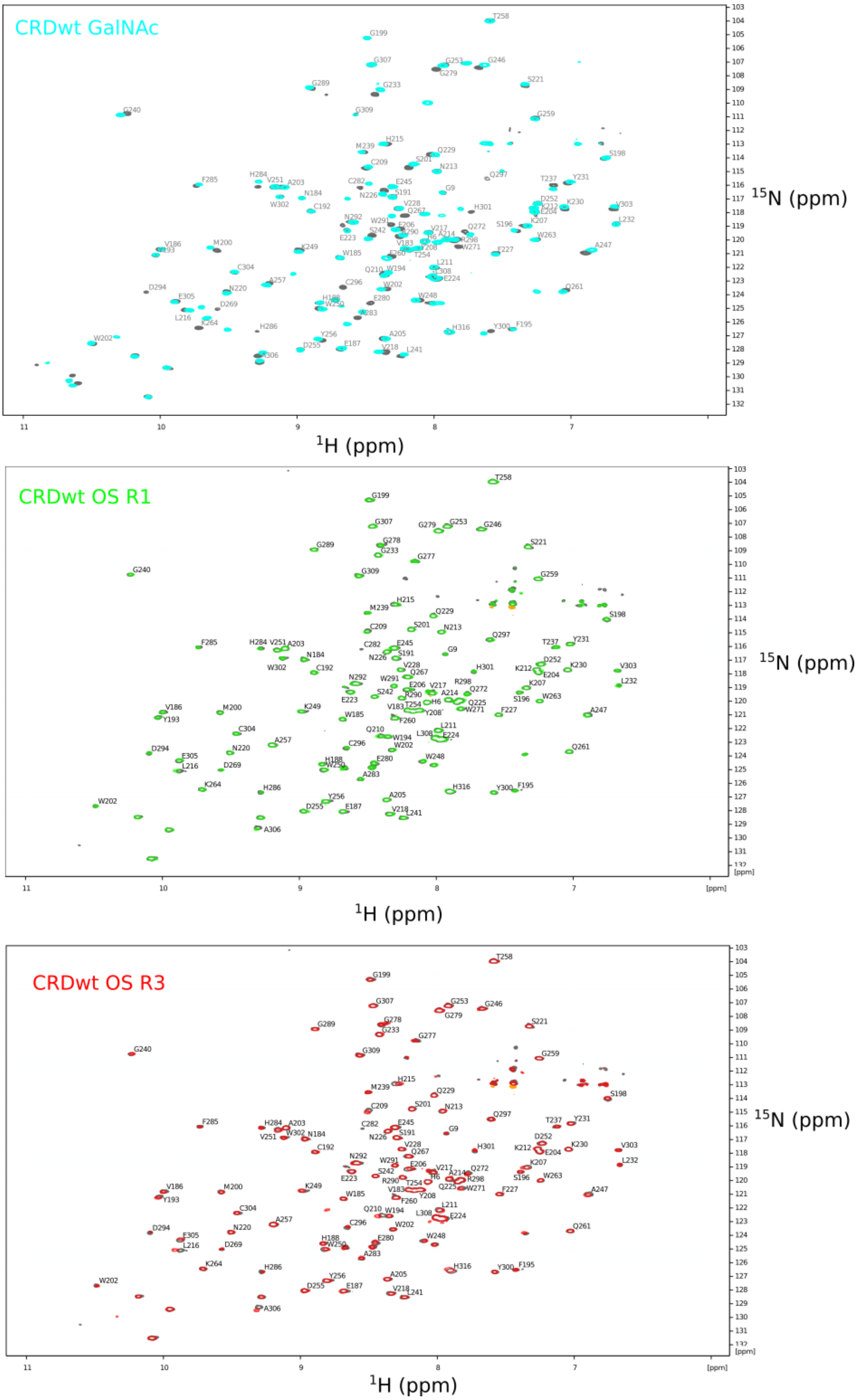


Figure S5. $^{15}N$-$^{1}H$ correlation spectra of MGL-CRD wt before(black) and after addition of 20:1 Glycan/protein ratio.

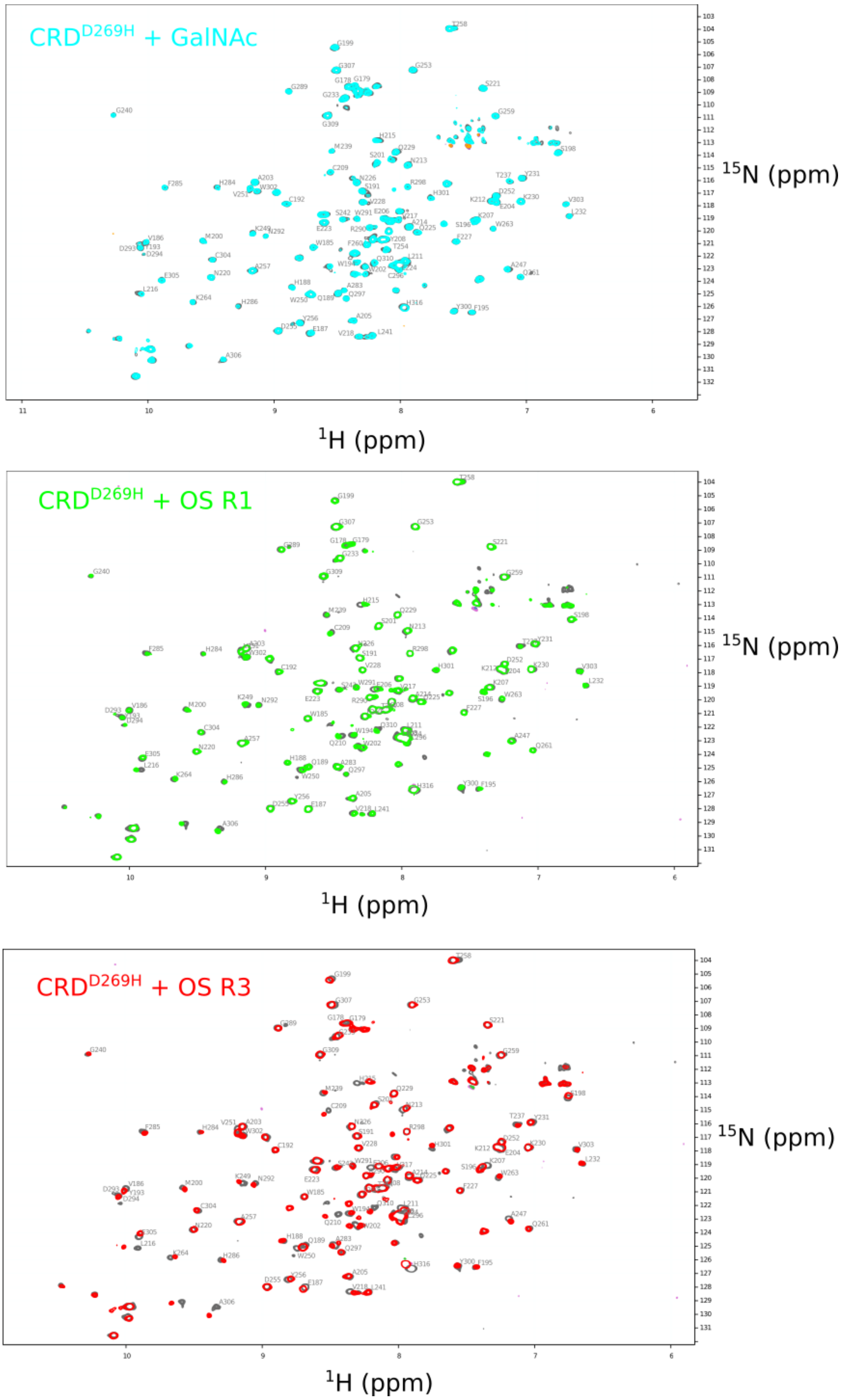


Figure S6. $^{15}N$-$^{1}H$ correlation spectra of MGL-CRD$^{D269H}$ before(black) and after addition of 20:1 Glycan/protein ratio.

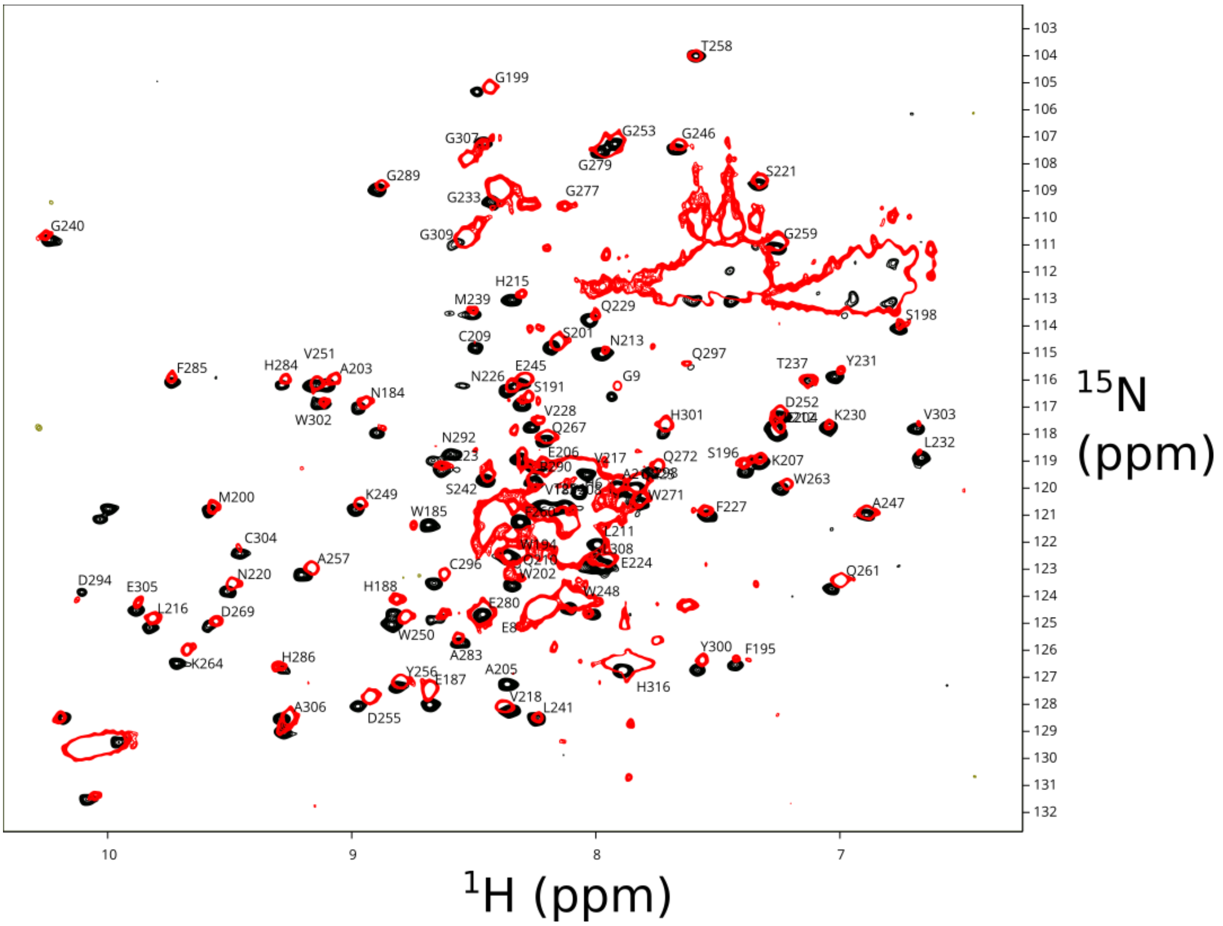


Figure S7 Overlay of $^{15}N$-$^{1}H$ correlation spectra of MGL-ECD (red) at 35°C and MGL-CRD(black) at 30°C. The assignments of the CRD that could be transferred by superimposition are indicated.

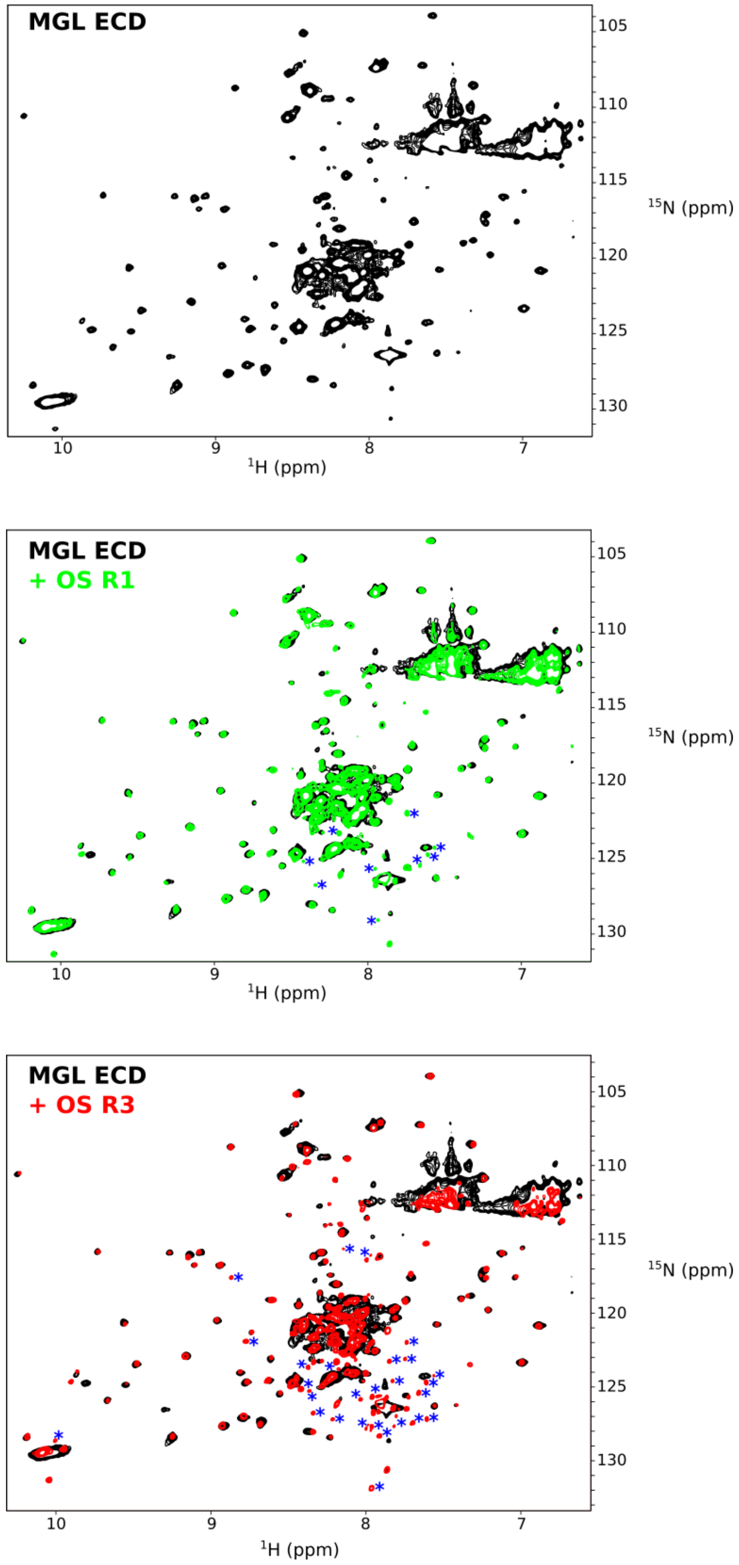


Figure S8. $^{15}$N-$^{1}$H correlation spectra of MGL-ECD wt before(black) and after addition of 2:1 Glycan/protein ratio. Blue asterisks correspond to new peaks appearing from minor protein degradation over time.

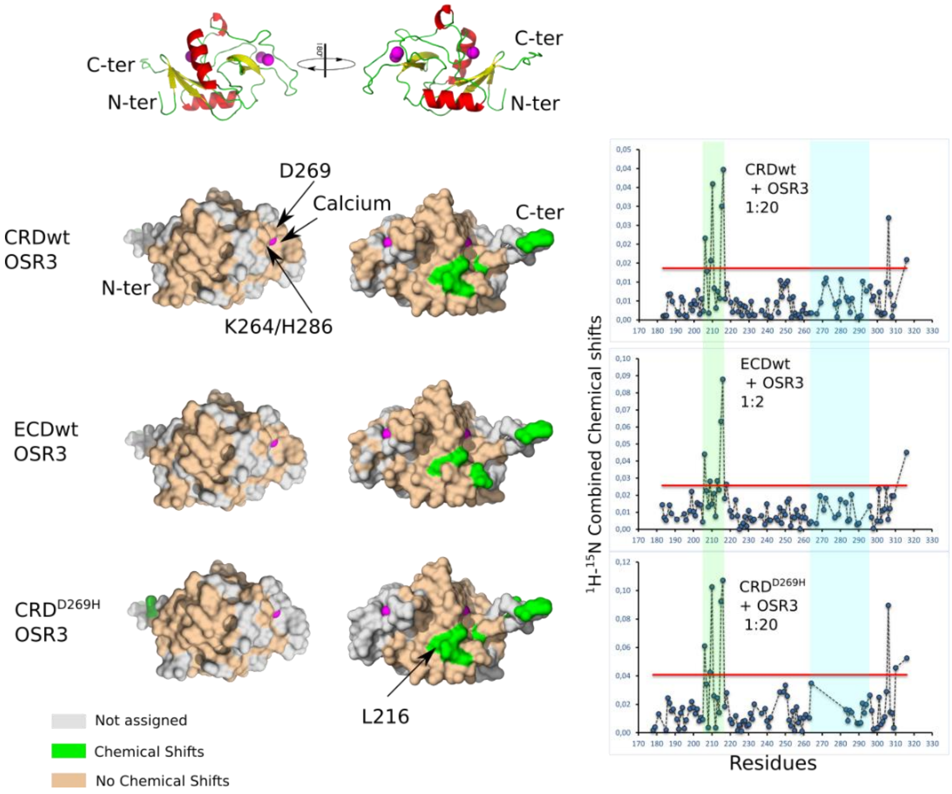


Fig S9. Chemical shifts Perturbations represented on MGL-CRD surface of MGL-CRD wt, D269H and ECD upon OS R3 interaction with the same color codes and orientations as in Fig 3. Histogram of CSP depending on residue number is shown on the right.

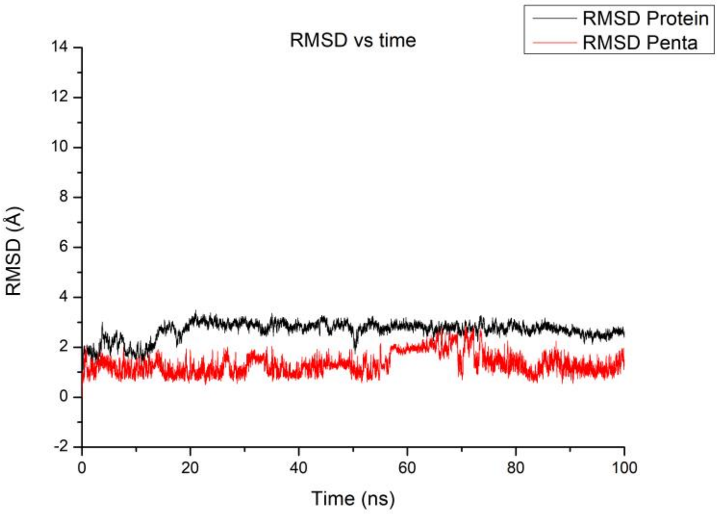


Fig S10: RMSD of the MGL (black) and OS R1 pentasaccharide (red) during the molecular dynamics calculated having the protein as reference.

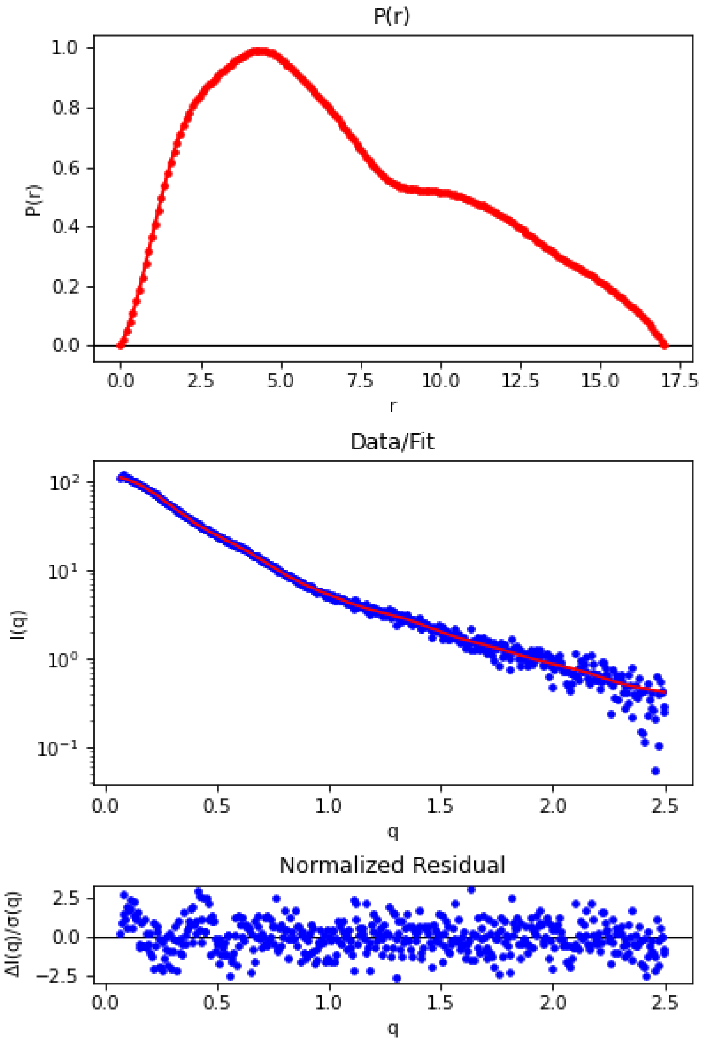


Fig S11: Pairwise distribution function used to calculate the SAXS envelope, with comparison (middle) with the experimental curve and associated residuals.

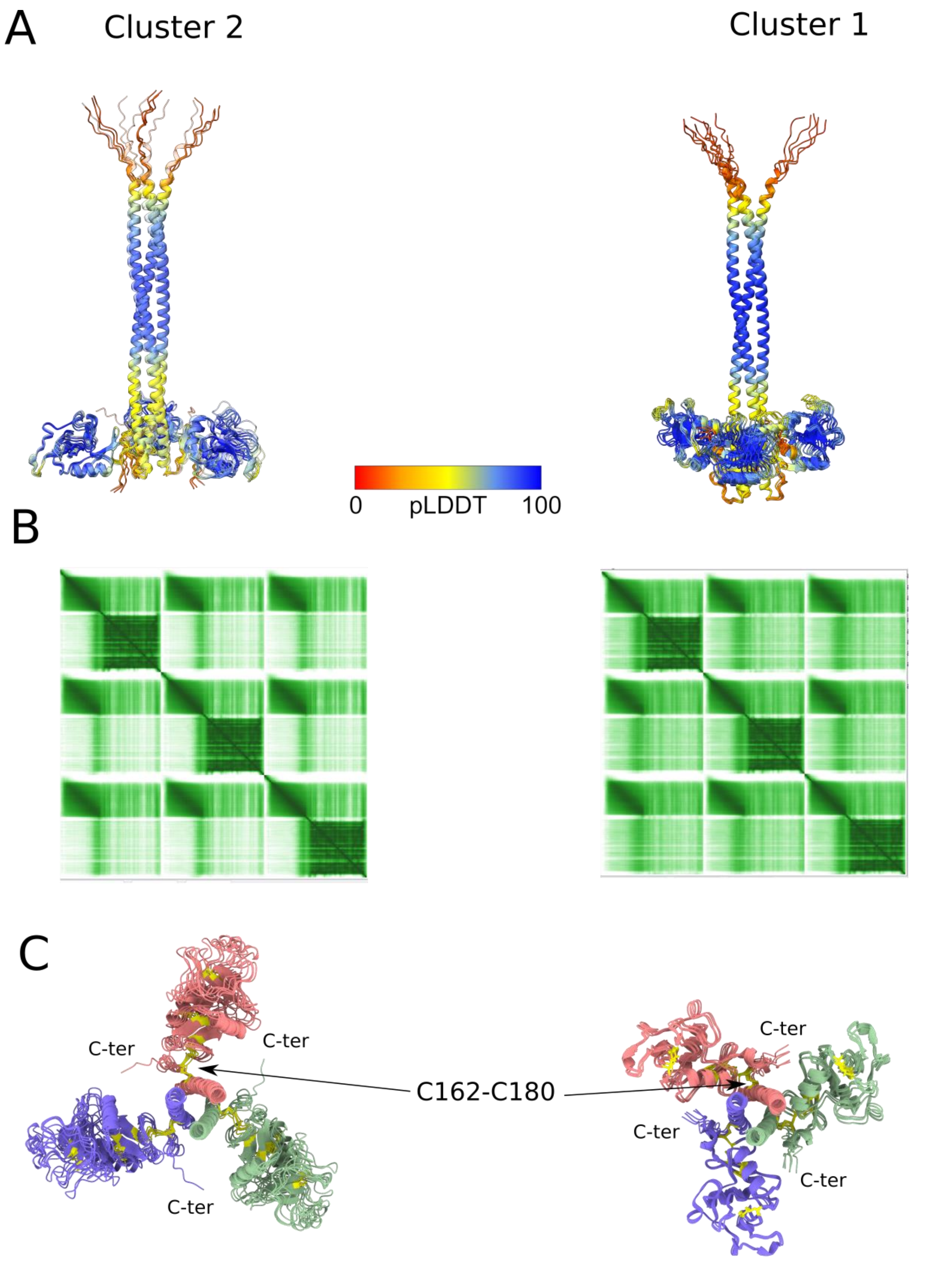


Fig S12 Alphafold models can be subdivided into two clusters. A) superimposed structures of both clusters with their PLDDT scores. The CRD structure is predicted with a 0.5 Å rmsd (across 128 residues 0.46 Å rmsd for cluster 1 and 0.52 Å for cluster 2) with respect to X-ray structure of MGL CRD (PDB:6PY1) B) The Prediction Alignment Error for representative structures of both clusters are showed and show little prediction of interactions between coiled-coil and CRD domains. C) structures of both clusters Superimposed on the coiled-coil motif. Disulfide bonds are shown in yellow.

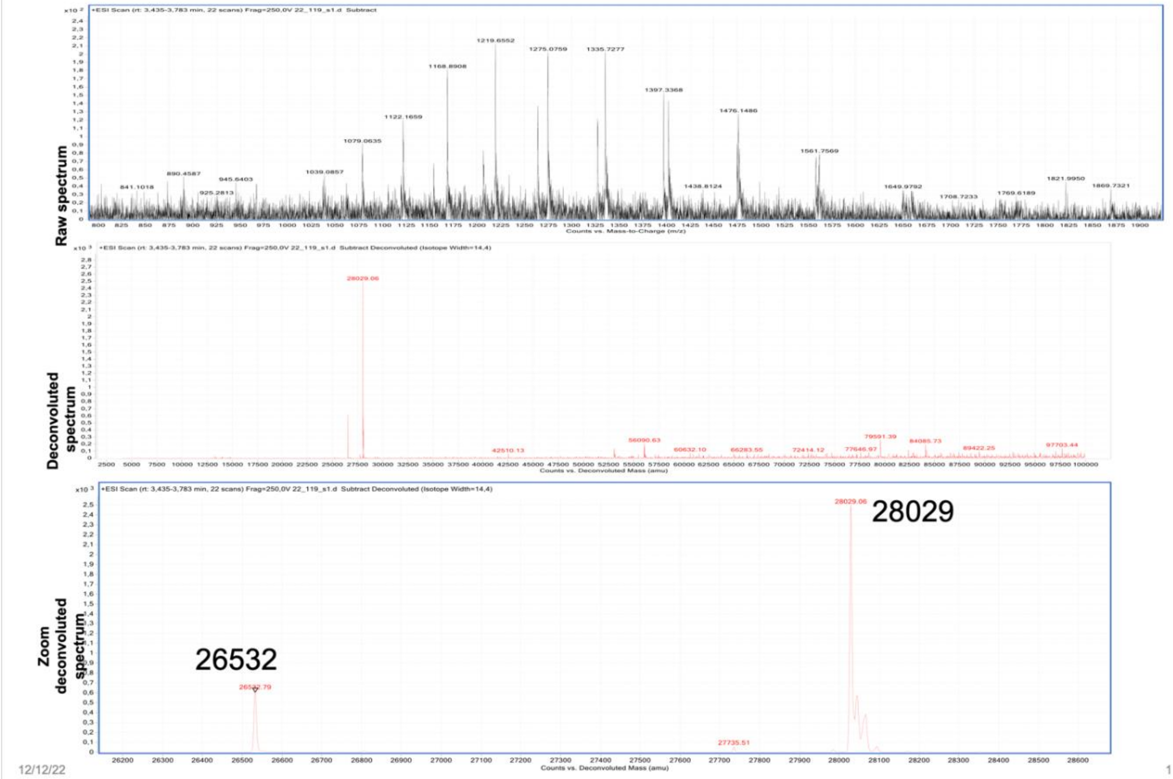


Fig S13. Electrospray Mass spectrometry of MGL-ECD. The expected mass of MGL-ECD is 28036.7 Da with loss of the first methionine. The main mass observed is 28029 Da corresponding to four oxidized cysteines. A minor peak at 26532 Da is probably a truncated form of MGL-ECD.

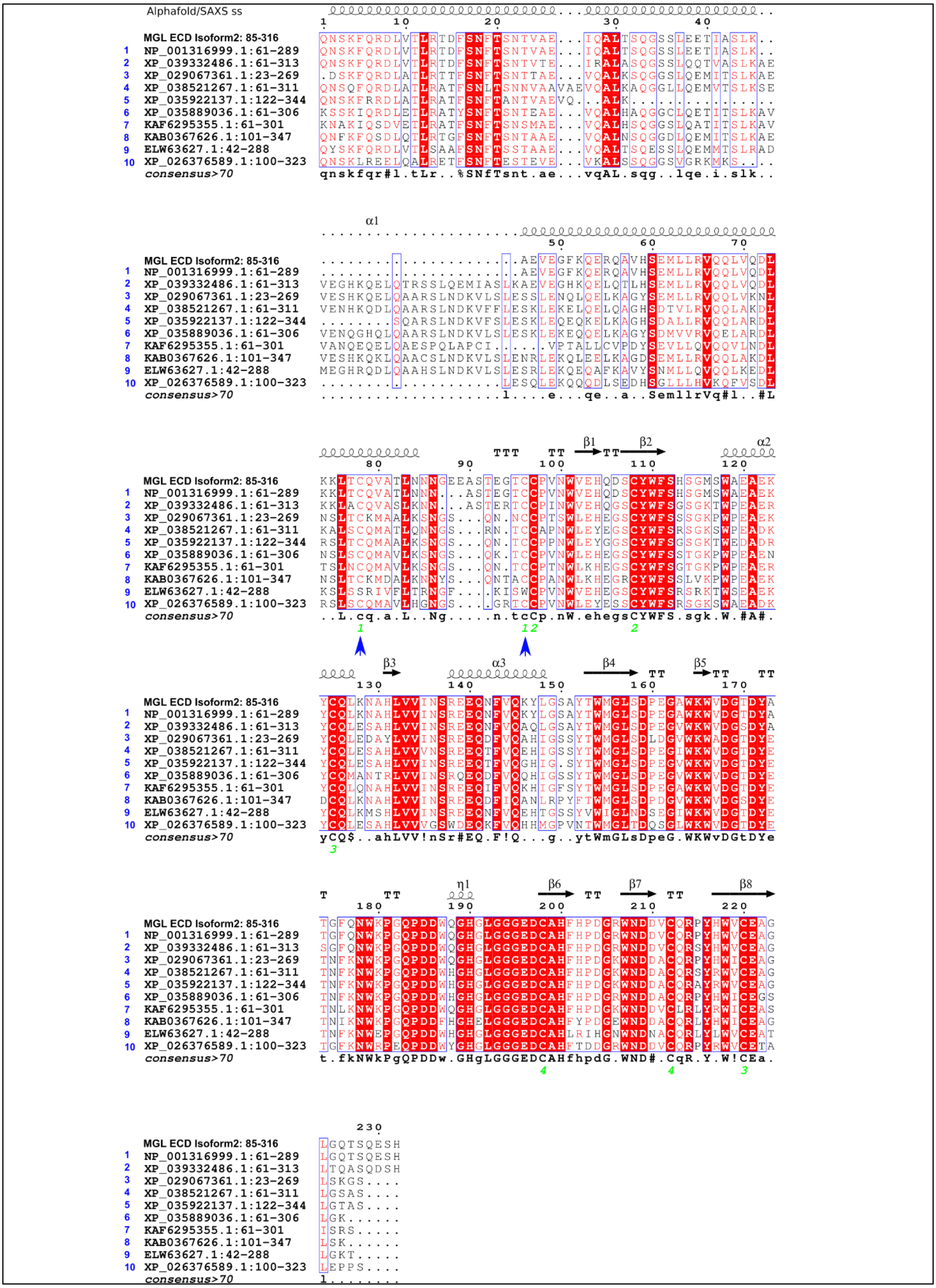

Figure S14: Sequence alignment of the MGL ECD sequence used in this study after a BLAST against clustered non-redundant database. Alignment was performed with clustalO and represented using Espript3 (Robert and Gouet, 2014). The secondary structure was extracted from the best alphafold/SAXS structure (ranked 5). Cysteines involved in disulfide bonds are shown with a number in green. Cys 162 and 180 are highlighted by a blue arrow.

| | Cluster Composition | Cluster Ancestor | Representative sequence | Query Cover | E value | % ident | Acc. Len |
|---|---|---|---|---|---|---|---|
| 1 | 14 member(s) 7 organism(s) | apes | CLEC 10 member A isoform 3 [Homo sapiens] | 100% | 1E-168 | 98.71 | 289 |
| 2 | 2 member(s) 2 organism(s) | primates | CLEC 10 member A isoform X1 [Saimiri boliviensis boliviensis] | 100% | 1E-140 | 77.34 | 313 |
| 3 | 2 member(s) 1 organism(s) | narwhal | CLEC 10 member A-like isoform X4 [Monodon monoceros] | 97% | 6E-119 | 65.74 | 269 |
| 4 | 6 member(s) 4 organism(s) | dog coyote wolf fox | CLEC 10 member A isoform X2 [Canis lupus familiaris] | 98% | 4e-113 | 63.92 | 311 |
| 5 | 2 member(s) 1 organism(s) | gray seal | ASGPR 1-like isoform X15 [Halichoerus grypus] | 98% | 4E-113 | 68.42 | 344 |
| 6 | 1 member(s) 1 organism(s) | pale spear-nosed bat | CLEC 10 member A-like isoform X1 [Phyllostomus discolor] | 97% | 9E-111 | 61.60 | 308 |
| 7 | 1 member(s) 1 organism(s) | bats | C-type lectin domain containing 10A [Myotis myotis] | 97% | 4E-108 | 62.45 | 302 |
| 8 | 1 member(s) 1 organism(s) | Reeves' muntjac | hypothetical protein FD755_020950 [Muntiacus reevesi] | 97% | 4E-103 | 61.20 | 349 |
| 9 | 1 member(s) 1 organism(s) | Chinese tree shrew | CLEC 10 member A [Tupaia chinensis] | 97% | 4E-99 | 60.56 | 509 |
| 10 | 1 member(s) 1 organism(s) | brown bear | ASGPR 1 isoform X1 [Ursus arctos] | 98% | 9E-97 | 60.09 | 323 |

Supplementary table 1: Cluster summary of sequences shown in the alignment on figure S14. For clarity, 1 cluster every 10 cluster was retained for the alignment, expect for sequence 5 which represents the closest ASGPR sequence. Clustered non-redundant database is the ncbi NR database clustered with each sequence within 90% identity and 90% length to other members of the cluster.

| Alphafold | chi2 SAXS | dockQ |
|---|---|---|
| 0 | 14,35 | 0,58 |
| 1 | 13,91 | 0,57 |
| 2 | 4,45 | 0,55 |
| 3 | 4,38 | 0,55 |
| 4 | 14,06 | 0,55 |
| 5 | 3,34 | 0,54 |
| 6 | 4,27 | 0,54 |
| 7 | 13,62 | 0,53 |
| 8 | 4,04 | 0,51 |
| 9 | 13,75 | 0,51 |

Supplementary Table 2: Alphafold models with their DockQ score and the $chi^2$ value with respect to the SAXS curve evaluated by crysol. Cluster 2 structures are colored in grey. The best matching structure is structure 5 with a $chi^2$ of 3.3 and the best from cluster 1, structure 7 ($chi^2$ 13.6)

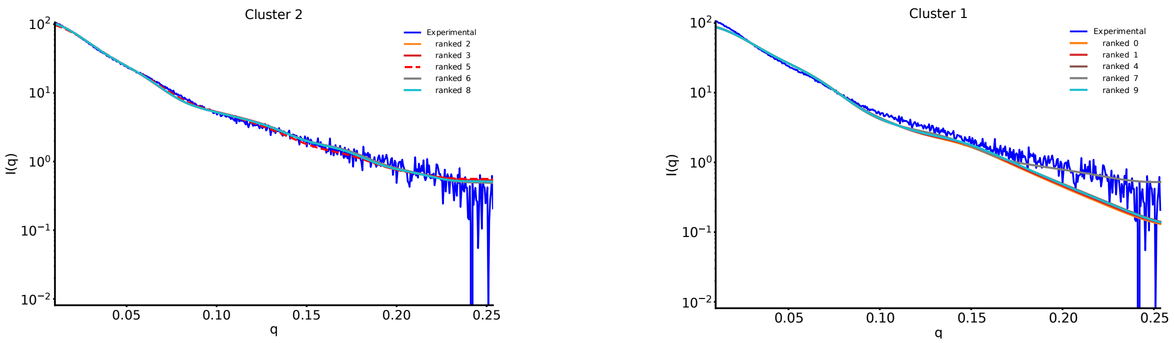


Fig S15 : Prediction of SAXS curves of the alphafold models and comparison with the experimental curve. $Chi^2$ values are in SI table 1.

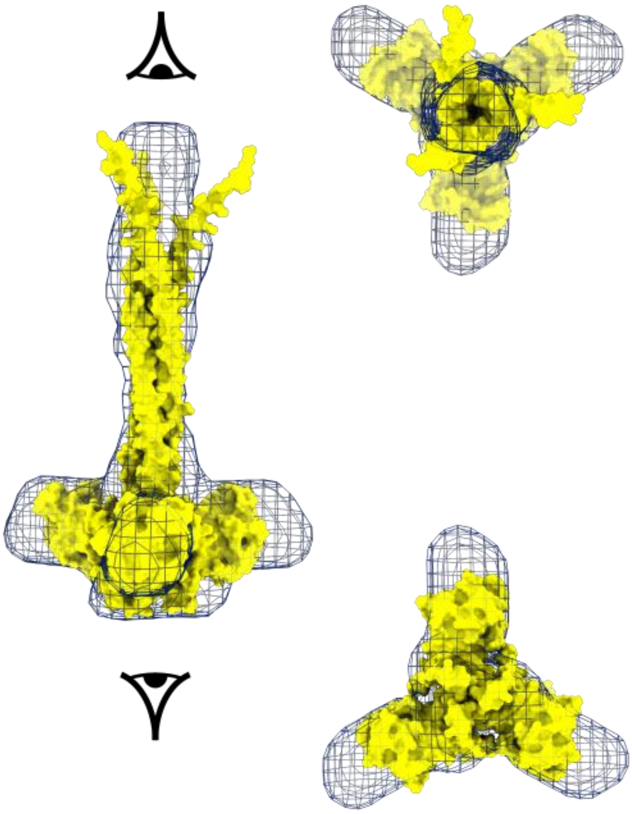

Fig S16 Comparison of cluster 1 structure with the best chi2 value with respect to the SAXS curve (ranked 7) is adjusted into SAXS calculated envelope.
